# Exploiting general independence criteria for network inference

**DOI:** 10.1101/138669

**Authors:** Petras Verbyla, Nina Desgranges, Sylvia Richardson, Lorenz Wernisch

## Abstract

Inference of networks representing dependency relationships is a key tool for understanding data derived from biological systems. It has been shown that nonlinear relationships and non-Gaussian noise aid detection of directions of functional dependencies. In this study we explore how far generalised independence criteria for statistical independence proposed in the literature are better suited to the inference of networks compared to standard independence criteria based on linear relationships and Gaussian noise. We compare such criteria within the framework of the PC algorithm, a popular network inference algorithm for directed acyclic dependency graphs. We also propose and evaluate a method to apply unconditional independence criteria to assess conditional independence and a method to simulate data with desired properties from experimental data. Our main finding is that a recently proposed criterion based on distance covariance performs well compared to other independence criteria in terms of error rates, speed of computation, and need of fine-tuning parameters when applied to experimental biological datasets.

## 1. Introduction

### 1.1. Motivation

Biological systems are driven by complex regulatory processes. In the analysis and reconstruction of such processes graphical models play a crucial role. Using network inference algorithms it is possible to derive regulatory models from high-throughput data, for example, from gene or protein expression data. A wide variety of network inference algorithms have been designed and implemented and necessitate common platforms for assessment, for example, the DREAM network inference challenges [11], to provide objective means for choosing reliable inference algorithms.

Inference algorithms are based on a variety of statistical principles. However, most rely on some form of estimating or testing the similarity or correlation between genes, for example, GeneNet [15] and MutRank [14], or on mutual information as does CLR [3] and RelNet [2], or on regression and feature selection as does MRNET [13] and Genie3 [9].

Using independence criteria for network inference have been suggested in contexts outside biological networks [10] and form the basis of the classic PC algorithm and its many variations [16]. Such algorithms exploit d-separation, the equivalent of statistical independence for graph structures, for the inference of directed acyclic dependency graphs (see for example, [16]).

Linear dependencies and Gaussian noise are typically assumed for most applications of the PC algorithm to continuous data. However, such assumptions are likely to be too restrictive in the case of many experimental datasets. Moreover, as a series of studies have shown (for example, [8]), the simplifying assumptions of linearity and Gaussian noise make it even more difficult to establish functional dependencies and their directions. They constitute limiting cases where for two dependent variables, for example, it becomes impossible to infer the direction of their functional dependency. These arguments strongly suggest that inference algorithms based on statistical independence should exploit nonlinear dependencies and non-Gaussian noise. The idea of combining the PC algorithm with a generalised independence criterion, the Hilbert-Schmidt Independence Criterion or HSIC, as independence oracle for conditional independence was proposed in [26] but not made operational.

The contribution of our study is threefold. First, we compare the performance of several independence criteria on biological experimental data. In particular, we compare the linear-Gaussian, the HSIC, and a further criterion based on distance, the Distance Covariance Criterion or DCC [24, 25], within the framework of the PC algorithm, when applied to protein expression data. Second, since not all criteria are available in a version that allows for testing conditional independence, we propose and test an approach that relies on residuals and requires only an unconditional version of an independence criterion. Third, the true network is rarely known when assessing algorithms. Hence, we also propose a simulation method that, starting from experimental data and a target network, produces simulated data according to the dependency structure of a target network but which are otherwise as close to the original data as possible in their noise characteristics and functional (possibly nonlinear) forms of dependencies. We demonstrate that such simulated dataset can be used successfully for differentiating between the performance of independence criteria for network inference. Finally, we make all algorithms and data available as a package for the R statistical environment [17].

We emphasise that the current study is not proposing a new inference algorithm and does not attempt to compare the performance of the PC algorithm to other network inference algorithms. Our aim is rather to compare and assess the relative merits of various independence criteria for network inference within the framework of a typical inference approach such as the PC algorithm and to explore problems and suggest solutions for their implementation and application to experimental data.

In the following sections we describe a representative dataset. We then provide some background on the inference of directed acyclic graphs using statistical independence as well as two generalised independence criteria. In the Results section their performance is compared on simulated as well as the original data.

### 1.2. Data

In order to assess general independence criteria for the inference of biological regulatory networks, we turn to a well studied dataset on protein expression from Sachs et al. [20]. The study comprises eight experimental datasets of single cell measurements. Each dataset reports the expression level of eleven proteins: RAF, MEK, ERK (aka P44.42), PLC*γ*, PIP2, PIP3, PKC, AKT, PKA, JNK, P38. The number of observations (cells) varies from 700 to 900 cells per dataset. Each dataset is characterised by the quantitative value of protein expression response to a specific stimulatory cue or an inhibitory intervention (listed in Table A1 in the supplementary material).

Protein expression levels were obtained by flow cytometry [20] measuring modification states of proteins, such as phosphorylation through antibodies. These are single cell measurements with each cell representing an independent observation. Only a few protein modifications are monitored. For example, PKC phosphorylates RAF at S497, S499, S259, however, only antibodies for RAF S259 were available. Consequently, some dependencies between protein states might be missed. Despite these shortcomings, some links between proteins are well established and shown in Figure 1 according to [20].

**Figure 1.:**
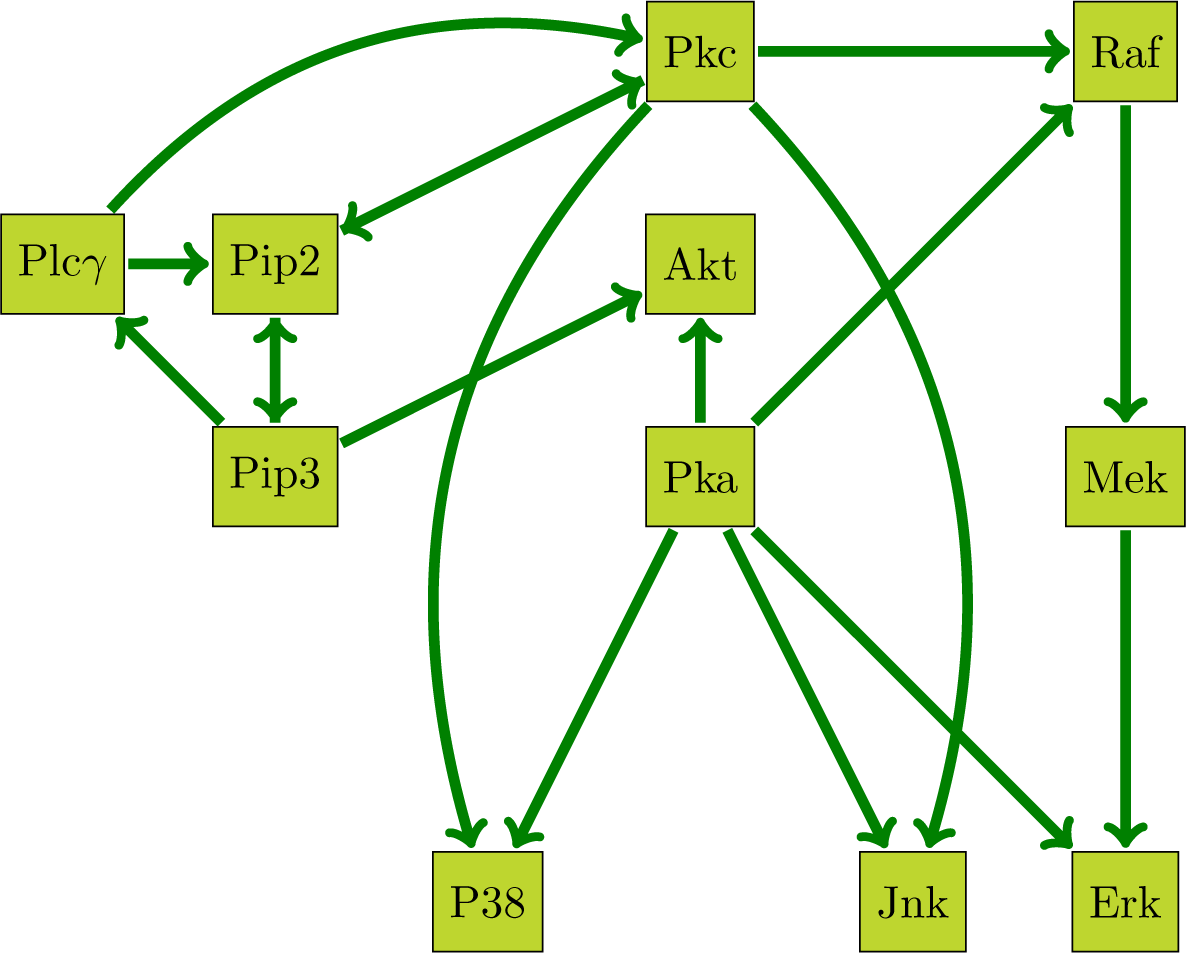
Summary of known dependencies (after [20]).

The graph in Figure 1 serves a twofold purpose. First, by exploiting the graph structure and by resampling the data, as described in more detail in the Results section, we generate datasets with characteristics close to real experimental datasets, but with known dependencies. They will serve as test sets for comparing the performance of inference algorithms. Second, using the original datasets, the graph provides a gold standard for measuring performance of algorithms on experimental data. Several limitations need to be kept in mind though. For some edges the direction of causal influence remains ambiguous. All inference algorithms considered here are based on the assumption that the dependency structure can be represented by a directed acyclic graph. Such assumption is likely to hold only approximately true in real datasets. Also the selection of proteins is by no means complete and it is likely that latent, unobserved variables induce additional dependencies. However, since we are primarily interested in a comparison of performances of independence criteria, these limitations are less problematic for the present study. There is no reason to belief that these inaccuracies in the knowledge of the true network should favor one approach unfairly over another. Conclusions about the relative merits of independence criteria based on the current datasets should generalise well to other data.

Figure A1 in the supplementary material shows boxplots for each variable and pairwise scatterplots between variables of dataset 8. Some dependencies are clear and close to linear, for example, RAF to MEK. Other dependencies are far less obvious, for example, RAF to PKC. This pattern of clear marginal dependencies between some of the related protein pairs but not all of them, is present in all eight datasets, as a reminder that network inference is not easily reducible to simple marginal correlation.

### 1.3. Probabilistic graphical models

We recall some terminology for probabilistic graphical models. For a full and comprehensive introduction see, for example, [23]. A graph **G** = (**V, E**) has vertices **V** and edges **E** ⊆ **V** *×* **V**. An edge between two nodes *V*_1_ and *V*_2_ can be either undirected, symbolically *V*_1_ − *V*_2_, or directed, symbolically *V*_1_ → *V*_2_. For a directed edge *V*_1_ *→ V*_2_, *V*_1_ is a parent of *V*_2_ and *V*_2_ is a child of *V*_1_. Two vertices connected by an edge or directed edge are adjacent. For a set of vertices **W** the set of all parents is denoted by Pa_**G**_(**W**). The degree *d*(*V*) of a node *V* is the number of nodes adjacent to *V*. A sequence of nodes (*V*_1_, …, *V*_*n*_), *V*_*i*_ *∈* **V**, forms a path if neighbouring nodes are connected. A directed path has all its edges directed in the same direction. A path is a cycle if (*V*_1_ = *V*_*n*_).

A *directed graph* contains only directed edges. It is a *directed acyclic graph* (DAG) if it contains no directed cycles. A graph is *undirected* if it contains only undirected edges and *mixed* if it contains both types of edges. A DAG is *compatible with a mixed graph* if the graphs agree on the directed edges. A *collider* (v-structure) is a triplet 〈*V*_1_, *V*_2_, *V*_3_〉 ⊂ **V** such that {*V*_1_, *V*_3_} ∈ Pa_**G**_(*V*_2_) and (*V*_1_, *V*_3_) *∉* **E**.

In a probabilistic graphical model the nodes **V** are associated with random variables with a joint probability distribution **P**. We will denote the random variables and the corresponding vertices by the same identifiers. In this study we are concerned with continuous variables and hence assume **P** is continuous and that it has a density *f*. For sets **X, Y, Z** of variables with conditional probability densities *f* (**X, Y** | **Z**), *f* (**X** | **Z**), and *f* (**Y** | **Z**), **X** and **Y** are conditionally independent given **Z** (denoted by **X** ⊥_*P*_ **Y** | **Z**), if *f* (**X, Y** | **Z**) = *f* (**X** | **Z**)*f* (**Y** | **Z**). A probability distribution **P** over the node set **V** is called *Markov* with respect to a DAG **G** if it permits the factorization

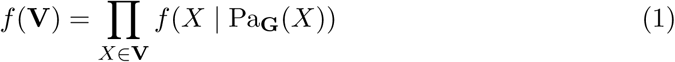

Symbolically, **X** ⊥_*G*_ **Y** | **Z** indicates that for a graph **G**, which is associated with a set of random variables **V**, and vertex sets **X, Y, Z** ∈ **V**, factorization (1) implies that **X** is independent of **Y** conditioned on **Z**. With this notation in place we see that **P** is Markov with respect to **G** if **X** ⊥_*G*_ **Y** | **Z** implies **X** ⊥_*P*_ **Y** | **Z** for all sets **X, Y**, and **Z**, that is, all independencies implied by **G** are in **P**. If, conversely, **X** ⊥_*P*_ **Y** | **Z** implies **X** ⊥_*G*_ **Y** | **Z**, that is, all independencies of **P** are implied by **G, P** is *faithful* to **G**. That is, if a distribution **P** is Markov and faithful with respect to a graph **G**, the graph represents all and only the independencies of **P** as captured in factorization (1). We will also say that **G** *represents* **P** *faithfully* in this case. Two DAGs are *Markov equivalent* if each distribution is Markov to either both or none of them. Consequently, in this case the two graphs are indiscernible based on probabilistic independency relationships alone.

### 1.4. PC algorithm

The PC algorithm [23] is a constraint based method for finding a DAG **G** that represents **P** faithfully. More precisely, if such graph exists, the algorithm returns a *partially directed graph* (PDAG), a mixed graph so that there exists at least one DAG which is compatible with the PDAG and represents **P** faithfully. The PDAG also has the property that for any undirected edge there exist two DAGs compatible with the PDAG, both representing **P** faithfully, but with the edge oriented in opposite directions. In this sense the PDAG is *maximally oriented*. If **P** can be represented faithfully by a DAG **G** and if there exists an independence oracle that returns, for each query triplet **X, Y, Z**, whether **X** ⊥_*P*_ **Y** | **Z** or not, [23] show that the PC algorithm returns a maximally oriented PDAG that is compatible with **G**.

The PC algorithm consists of three distinct phases. The first (skeleton) phase finds the skeleton of the PDAG, that is, it finds all adjacencies based on an independence oracle. The second (collider) phase finds all colliders (*V*_*i*_, *V*_*k*_, *V*_*j*_) and directs edges *V*_*i*_ → *V*_*k*_ and *V*_*j*_ → *V*_*k*_. The third (transitive) phase applies the Meek rules [12] to extend all directions found in the collider phase to the rest of the PDAG ([12, 26] give a succinct description of details).

In addition, the PDAG can be extended using background knowledge about the direction of some of its undirected edges. If the background knowledge is compatible with the underlying distribution **P**, iterative application of an extended set of Meek rules results in an a PDAG-K (PDAG with background *knowledge*) that again is maximally oriented in the sense that for each undirected edge there exist two DAGs compatible with the PDGA-K and both representing **P** faithfully as well as agreeing with the background knowledge, but which contain opposite directions of the edge (see [12] for details).

In practical applications with finite samples an independence oracle is usually unavailable and is replaced by a statistical test of independence. Throughout the paper we use tests (described in detail in the next section) that allow us to reject the null hypothesis of independence in terms of *p*-values, that is, low *p*-values indicate dependencies. A consequence of sampling error is that the PC algorithm might not produce the correct PDAG, but might instead produce a graph with under or over predicted edges, wrong directions of edges, doubly directed edges directed in both directions, and even cycles. In most circumstances it is appropriate for the algorithm to return such ambiguous graph and leave it to the user to decide which edges or directions to ignore.

A fixed cutoff level *α* is used for all decisions on independence in the PC algorithm. Since edges indicate dependencies, higher levels of *α* usually result in the acceptance of more edges and denser graphs.

#### 1.4.1. Additive Noise model

Tests for conditional independence are based on certain assumptions. A popular one is that dependencies between a child and its parents can be modelled by a linear function with additive independent Gaussian noise. Unfortunately, apart from being rarely fulfilled in practise, these assumptions make it actually more difficult to identify directions of influence between variables. Several authors, for example, [8] or [26], therefore assume more general noise models. In *additive noise models* the dependency of a variable on its parents is modelled by a nonlinear function and additive noise (not necessarily Gaussian), that is, *V*_*i*_ = *f*_*i*_(Pa_**G**_(**V_i_**)) + *є*_*i*_, with *є*_*i*_ independent for each *i*.

If data are generated by such process the assumption of an additive noise model, which includes the linear Gaussian noise case, has a twofold advantage. First, independence tests used in the PC algorithm based on nonlinearity and general noise distributions are more accurate than tests based on assumptions of linearity or Gaussian noise distributions resulting in a more accurate PDAG. Second, exploiting the inherent improvement in establishing the correct direction for undirected edges, more edges can be directed than through consideration of independence relationships alone (as in the PDAG). Meek’s rules [12] for background knowledge can then be applied to obtain a PDAG-K to direct additional edges of the PDAG. This is achieved in the *generalised transitive phase* algorithm in [26] who show that it results in maximally directed mixed graphs for a slightly wider class of models than just additive noise models.

## 2. Methods

Our aim is to employ the PC algorithm and to apply a generalised transitive orientation phase to obtain a PDAG-K representing a probability distribution according to the additive noise model. For this purpose we need to specify an independence oracle that is suitable for nonlinear relationships and non-Gaussian noise. In the following we provide a summary of two criteria, the *Hilbert-Schmidt Independence Criterion* or HSIC and the *Distance Covariance Criterion* or DCC, and describe our implementations.

[21] show that the DCC is actually a version of the HSIC for a specific kernel. However, here we apply the HSIC with the widely used squared exponential kernel as explained in the following. In a way, comparing the HSIC with the DCC in this study is comparing the HSIC with two very different kernels, one motivated by a popular choice of the kernel function, the other rather indirectly via a distance correlation approach. However, the computational requirements and implementation issues of HSIC (with a squared exponential kernel) and of DCC are very different, as we discuss now.

We assume we have *n* samples *v*_*i*_ from a set of variables **V** sampled from a distribution **P**. We are interested in establishing independence of subsets of variables **X** *⊆* **V** and **Y** *⊆* **V** or their independence conditioned on variables **Z** *⊆* **V**. We denote the measurements corresponding to **X, Y**, and **Z** for sample *i* by *x*_*i*_, *y*_*i*_, and *z*_*i*_.

As outlined above the PC algorithm requires an independence oracle that states whether **X** *⊥*_*P*_ **Y** or **X** *⊥*_*P*_ **Y** | **Z** based on samples *v*_1_, …, *v*_*n*_. [26] suggest using the HSIC as independence oracle for the PC algorithm. In the following we will compare the performance of the two independence criteria, the *Hilbert-Schmidt Independence Criterion* or HSIC and the *Distance Covariance Criterion* or DCC. We call the PC algorithm based on HSIC, following [26], *kernel-PC* or kPC, and, in analogy, the PC algorithm based on DCC, *distance-PC* or dPC. We will define further variants of the kPC below.

### 2.1. Hilbert-Schmidt Independence Criterion

For a comprehensive introduction to the HSIC see for example [22] or [4]. For our purposes it is sufficient to describe the calculation of the HSIC statistic for a finite sample {(*x*_1_, *y*_1_), …, (*x*_*n*_, *y*_*n*_)}. The HSIC is based on a *kernel* function, a similarity function between sample points. As kernel function we use a Gaussian kernel 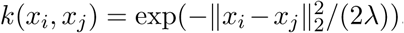, where the kernel width *λ* is a parameter that needs to be carefully selected (see the Results section). Let *K* and *L* be the Gram matrices associated with kernel functions *k* and *l*, that is *K*_*i,j*_ = *k*(*x*_*i*_, *x*_*j*_) and *L*_*i,j*_ = *l*(*y*_*i*_, *y*_*j*_). The centred Gram matrices are 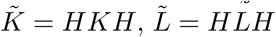, where 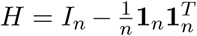 (here I_*n*_ is the *n*-dimensional identity matrix and **1**_*n*_ is a vector of ones of length *n*). An estimate 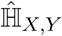 of the HSIC is then given by

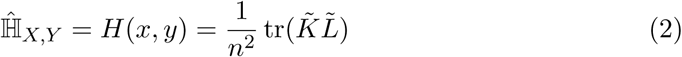

where tr(*A*) is the trace (sum of diagonal elements) of a matrix *A*. Generally, *H* (*x, y*) is close to 0 when *X* and *Y* are independent. [4] also provide an estimator for a *conditional* version 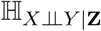 of the HSIC for a sample set (*x*_*i*_, *y*_*i*_, *z*_*i*_), *i* = 1, …, *n* as

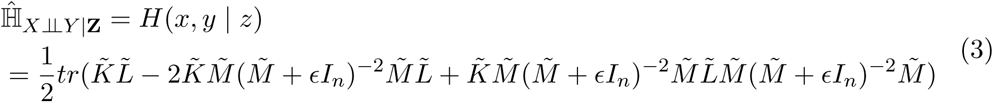

where 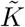, and 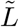 are defined as above for *x*, and *y*, and 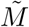 is the analogous Gram matrix for z. ∈ is a regularization parameter that needs to be carefully selected (see the Results section). The calculation of 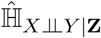 is computationally very expensive, but some simplifications are introduced in [26].

If we use the PC algorithm with the HSIC as independence oracle we obtain algorithm kPC.

### 2.2. Tests of (unconditional) independence for kPC

[5] suggest two ways of calculating a *p*-value for the HSIC statistic (2), a permutation test and a test based on an approximation using the Gamma distribution.

**Permutation test**. The first test is a simple permutation test where *r* permutations of the form 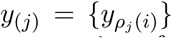, *j* = 1, *…, r*, for permutations *ρ*_*j*_ of sample indices are created. The proportion of permutations *ρ*_*j*_ for which the HSIC estimator (2) is larger than the HSIC of the original dataset, that is *H* (*x, y*_(*j*)_) *> H* (*x, y*), is an estimate of the *p*-value for rejecting the null hypothesis of independence. The underlying assumption is that permuting *y* removes any dependency between *x* and *y*.

Computing *H* (*x, y*) is expensive with complexity *O*(*n*^3^), *n* the sample size. [26] suggested an incomplete Cholesky factorization with *m* steps to reduce the complexity to *O*(*nm*^3^) for a chosen *m < n*. That is, *K* and *L* are approximated by 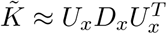, and the matrix of eigenvalues *D*_*x*_ of size *n × m* and *m × m*, respectively. Similarly 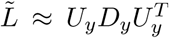. This factorization is suitable since, with quickly decaying kernel functions, Gram matrices often have a low effective rank. Instead of permuting the entries of *y* and recalculating the Gram matrix, we exploit the one-to-one relationship between samples *y*_*i*_ and the rows and columns of the Gram matrix *L*, since *L*(*i, j*) = *l*(*y*_*i*_, *y*_*j*_) (with a symmetric kernel function *l*). In the calculation of *H* (*x, y*_(*i*)_) we use 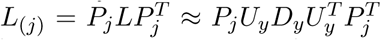, with a permutation matrix *P*_*j*_ permuting rows according to permutation *ρ*_*j*_. This means that an incomplete Cholesky decomposition needs to be performed only once for all permuted datasets *y*_(*i*)_, and consequent values *H* (*x, y*_(*i*)_) can be obtained simply by permuting coordinates of the eigenvectors in *U*_*y*_ (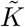 is kept the same).

**Gamma Test**. The value of the asymptotic distribution of the empirical estimate *H* (*x, y*) of the HSIC under the null hypothesis of independence is approximated by a Gamma distribution: *H* (*x, y*) *∼* Gam(*α, θ*) where *α* is the shape parameter and *θ* is the scale parameter calculated as

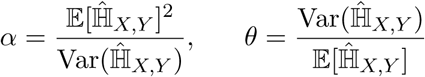

To compute this distribution we use estimates of the mean and the variance from sample points under the null hypothesis as provided by theorems 3 and 4 in [5]. The *p*-value is then obtained as upper-tail quantile of *H* (*x, y*).

### 2.3. Test of conditional independence for kPC

In the PC algorithm an oracle for conditional independence **X** ⊥_*P*_ **Y** | **Z** is required. We explore two approaches. The first, *permutation-cluster test* suggested by [26], is based on a conditional version of HSIC from [4]. The alternative test we propose here is based on *residuals*. It is simpler in that it only requires an unconditional version of the HSIC and can be readily applied to other independence criteria for which there is no conditional version readily available that allows integration of a conditioning set of variables, as in the case of the DCC.

**Permutation-cluster test**. As suggested in [26], in order to obtain a *p*-value for rejecting independence based on the estimator (3) for conditional HSIC criterion, the samples are clustered according to the Euclidean distance between the *z* coordinates of samples. Sample labels of *y* are only permuted within each cluster, thus ensuring that the permuted samples break dependency between *x* and *y* for an approximately fixed *z* but retain their dependency on *z*. For the clustering we use a k-means algorithm [[6]] (R function kmeans()). A larger number of clusters is desirable to achieve an almost constant *z* within each cluster. On the other hand, enough samples are required in each cluster to achieve a permutation of labels that breaks any conditional dependency between *x* and *y*. For the sample sizes considered here, good results were obtained with a constant cluster number of 10.

**Residuals test**. As a simpler alternative to obtain *p*-values for the conditional HSIC we propose to test residuals for independence based on any unconditional test of independence. The residuals *r*_*x*_ and *r*_*y*_ are obtained by regressing *x* and *y* on *z* in a nonlinear fashion. The regression removes the dependencies between *x* and *y* due to *z* and consequently the residuals should be independent if **X** ⊥_*P*_ **Y** | **Z**. For regression we use a generalized additive model (GAM, see [7]) as implemented in the R function `gam()` in the library mgcv [27, 28] with default settings). That is, we regress *y* on a set of variables 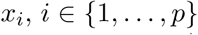 as 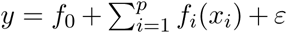 where *f*_*i*_ are spline functions (selected by cross-validation) and *ε* is Gaussian noise. We have now the option of using either the permutation or Gamma test of section 2.2 on the residuals.

### 2.4. Distance covariance

The distance covariance has been suggested as an alternative measure of independence to the HSIC by [24] and [25]. An estimate of the distance covariance for a set of *n* samples {(*x*_1_, *y*_1_), …, (*x*_*n*_, *y*_*n*_)} is obtained as follows. For variable *X* we define

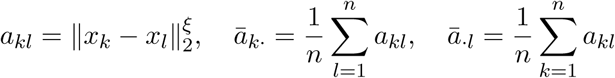

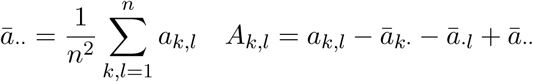

with the power parameter ξ. Similarly, we define

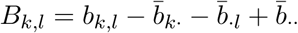

for the variable *Y*. An estimator for the distance covariance is then obtained as

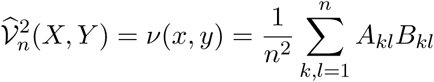

If we use the PC algorithm with the DCC as independence oracle we obtain algorithm dPC.

### 2.5. Test of (unconditional) independence for dPC

The independence test is already implemented by `R` in the `energy` package [19] with the function `dcov.test()`. The test is implemented as a permutation test. The output *p*-value is calculated as 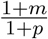 where *m* is the number of replicates which are greater than the observed value of the statistic dCov(X,Y) and *p* is the total number of replicates (personal communication with M.L.Rizzo).

### 2.6. Test of conditional independence for dPC

We generalize the test of the previous Section 2.5 to a conditional version by applying the (unconditional) independence test to residuals formed as in section Residuals test of 2.3.

## 3. Results

We first investigate the effectiveness of the independence criteria in finding dependencies in small simulated examples and explore parameter settings. We continue with a larger scale example with data simulated by permutation resampling from real data. Finally we present our results for the datasets from [20].

### 3.1. Testing unconditional independence criteria

We expect the effectiveness of the independence criteria to depend crucially on the signal to noise ratio. We therefore tested the HSIC and DCC on 300 samples simulated from 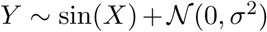, 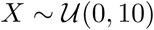 for varying noise levels *σ*^2^ and signal range of 2 from *−*1 to 1. *X* and *Y* are dependent, hence independence should be rejected with low *p*-values. The simulated data are shown in the figure A2 in the supplementary material. Table 1 lists *p*-values for combinations of methods from Sections 2.2 and 2.5 and varying noise levels *σ*. All the *p*-values are a mean of 100 replications of the test. The size of the simulated sample is 300, as this is a reasonably typical sample size for high-throughput experiments. At *σ* = 10 variables *X* and *Y* are effectively independent. As expected the *p*-value is less and less reliable for detecting dependency for samples with increasing noise levels. In this simple test both criteria, HSIC and DCC, behave similarly.

**Table 1.:** Testing independence criteria. All the *p*-value estimates are the mean of 100 *p*-values from repetitions for each of the three tests. The size of the simulated sample is 300. The DCC and HSIC *p*-values are obtained from 1000 permutations each.

| Test | $\sigma = 1$ | $\sigma = 2$ | $\sigma = 5$ | $\sigma = 10$ |
| --- | --- | --- | --- | --- |
| HSIC permutation | < 0.001 | 0.04 | 0.37 | 0.48 |
| HSIC gamma | 4e-13 | 0.04 | 0.38 | 0.49 |
| DCC | < 0.001 | 0.06 | 0.37 | 0.47 |

The HSIC depends on a kernel width parameter *λ*. Figure. 2a) to c) show *p*-values for different HSIC tests when the kernel width *λ* varies from 0.001 to 1000. We note that in order to reject independence successfully *λ* needs to be chosen carefully, particularly with higher noise. If *λ* is very small, then *k*(*x, y*) *≈* 0 for almost all *x* ≠ *y*, and the Gram matrix is close to the identity matrix. If *λ* is too large, then *k*(*x, y*) *≈* 1, for all *x, y* and the Gram matrix is ill-conditioned. In either case any dependency variables is hard to detect. Based on these figures we choose a kernel width in the range *λ* ∈ (0.5, 9). Furthermore we observe that the permutation and Gamma tests both with and without incomplete Cholesky decomposition always give very similar results for this range of *λ*. Therefore, for further analysis we will use the Gamma test with an incomplete Cholesky decomposition since it is computationally most efficient (as seen in Figure A4 of the supplementary material).

**Figure 2.:**
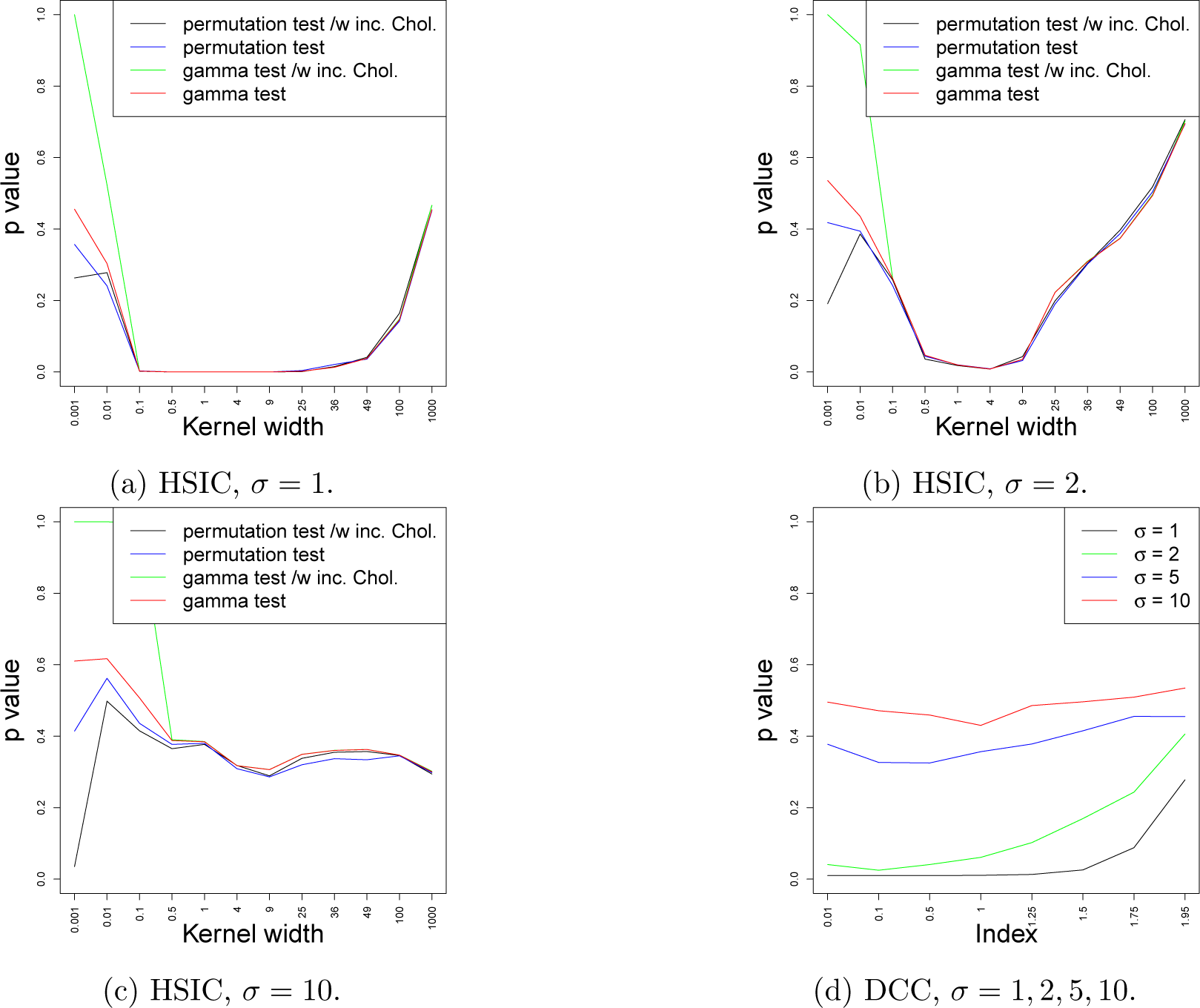
Dependency of *p*-values of HSIC and DCC tests on varying parameters, kernel width *λ* for HSIC and index *ξ* for DCCC.

Figure 2d) shows the dependency of *p*-values on the power parameter ξ of the DCC. For simplicity we set ξ = 1, although smaller values might work even better.

### 3.2. Comparison of network inference algorithms

The performances of the algorithms are compared using ROC curves sensitivity over specificity while varying the *p*-value cutoff required by the oracle for statistical independence in the PC algorithm. Unless otherwise stated we will focus on the absence and presence of edges in the inferred graphs ignoring their direction when calculating specificity and sensitivity.

Three types of datasets are considered: data simulated from a simple known network, data obtained by permuting residuals after fitting a network to data from [20], and finally the original data from the same study. Parameters of the algorithms were fixed as in Table 2.

**Table 2.:** Free parameters for kPCs and dPC.

| Parameter | kPC-Clust | kPC-Resid | dPC |
| --- | --- | --- | --- |
| # of Permutations | 300 | 300 | 500 |
| Kernel width $\lambda$ | 1 | 1 | NA |
| Regularization parameter $\epsilon$ | 0.1 | NA | NA |
| # of clusters | 30 | NA | NA |

For easy reference we label the algorithms as follows. The standard PC algorithm as implemented in the `R` package `pcalg` with the `gaussCItest` criterion (implementing Fisher’s *z*-test for correlation) is labelled PC. The PC algorithm based on the DCC from Section 2.4 is labelled dPC. The PC algorithm based on the HSIC from Section 2.1 is labelled kPC. The kPC version based on the Permutation-cluster test of Section 2.3 is labeled kPC-Clust. The kPC version based on the Residuals test of Section 2.3 with the Gamma approximation is labelled kPC-Resid.

#### 3.2.1. Data simulated from artificial network

The relationships between the nodes are described in Figure 3b. Since the network contains nonlinear relationships and non-Gaussian noise, as expected, the PC algorithm performed worst. Close to perfect performance was achieved by dPC, kPC-Clust and kPC-Resid. Figure A3a shows the spread of ROC curves when simulating data repeatedly. dPC, kPC-Resid, and kPC-Clust outperformed PC in 100, 98, and 100 out of 100 cases, respectively.

**Figure 3.:**
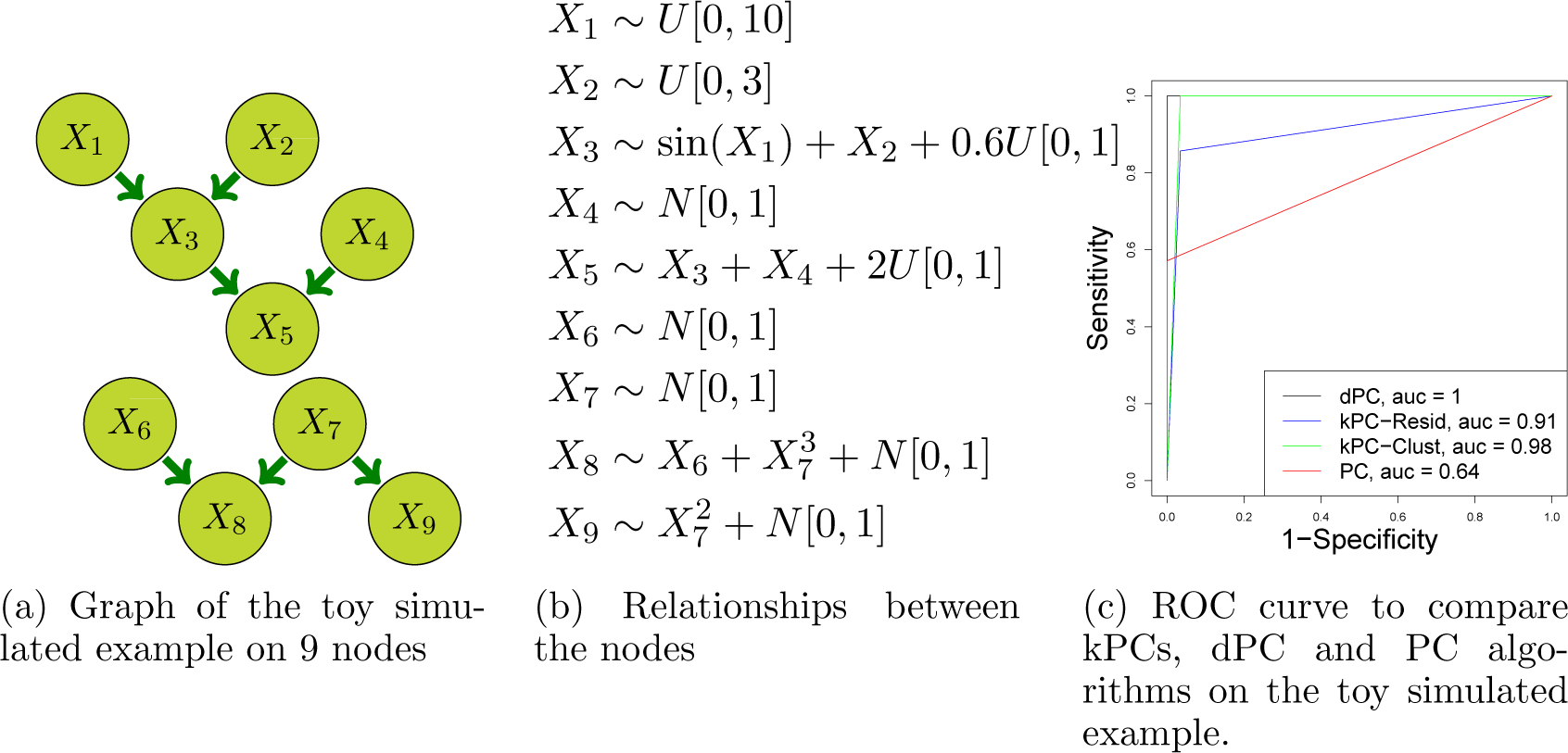
Toy simulated example on 9 nodes and 300 observations.

#### 3.2.2. Data simulated by resampling

Next we compared the performance of the algorithms on samples obtained by fitting a plausible network to data in [20] and resampling residuals. In this way we still control the structure of the underlying network but obtain more realistic noise distributions.

As outlined in the introduction, the data consist of eight datasets of expression levels of eleven proteins, each dataset obtained after specific experimental interventions. Here we present only the results from the dataset 7, the rest is provided in the supplementary material in Figure A5. Protein PKC was inhibited for dataset 7. Since PKC was externally modified no causal arcs lead into PKC. The network is that of Figure 1 with arcs into PKC removed.

For the simulation we used the causal model of Figure 4a. The data generation starts from parentless nodes (PKC and PKA). These variables are assigned the original values from the samples in the experimental dataset. Next, recursively iterating over nodes whose parents already have assigned values, a generalised additive model similar to section 2.3 is fitted to obtain mean estimates and residuals for the experimental sample values of the focus node when regressing on the values previously assigned to its parents. These residuals are permuted before being added to mean estimates to obtain resampled values to assign to the focus node.

**Figure 4.:**
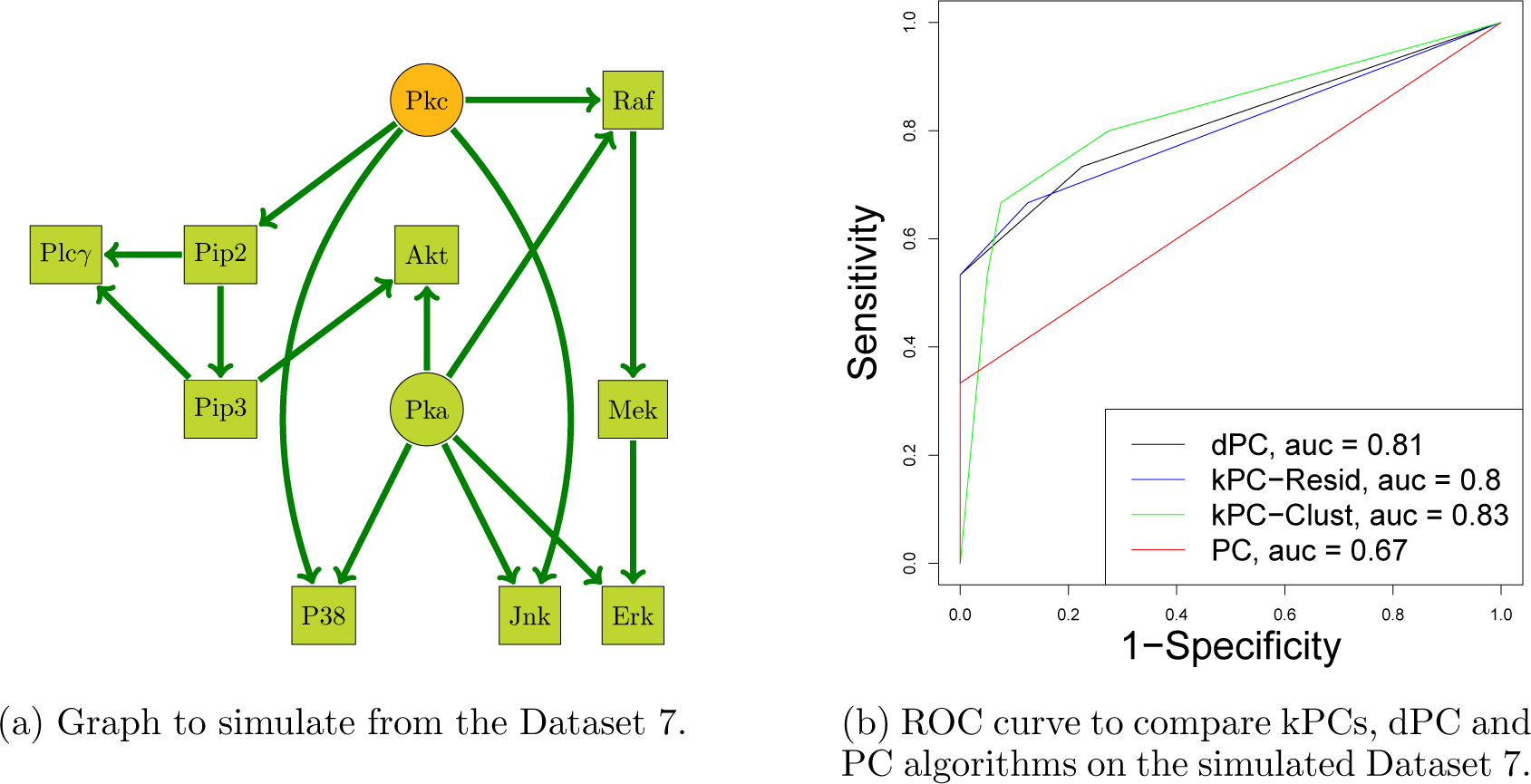
Data simulated from the Dataset 7, with non-reduced noise.

This procedure ensures that only the assumed dependencies as captured in the nonlinear regressions on parents are maintained, while all other dependencies are removed by permuting residuals. On the other hand, the noise characteristics of the original data are maintained to some degree. In particular, some focus nodes show little functional dependence on their parents, that is, a very low signal to noise ratio. This is a characteristic of experimental data as well. In order to explore the influence of this signal to noise ratio, additional data sets are simulated with residuals scaled down by a factor *k* before being added to mean estimates, improving on the signal to noise ratio.

Figure 4b shows that all the PC versions based on general independence criteria significantly outperform the traditional PC algorithm. dPC, kPC-Resid and kPC-Clust result in areas under the ROC curve of greater than 0.8 while that of PC is only 0.67. Performance is worse than for the toy example above. This is mainly due to a small signal to noise ratio for many relationships: on visual inspection many relationships in Figure 1 are hardly noticeable in the data. This results in the regression step not capturing much signal. On the other hand, real data are likely to show this type of noise characteristics. The effect of varying the scaling factor *k* for the residual noise is shown in Figure 5. Generally, as expected, with lower noise performance improves. The dPC version is responding well to lower noise.

**Figure 5.:**
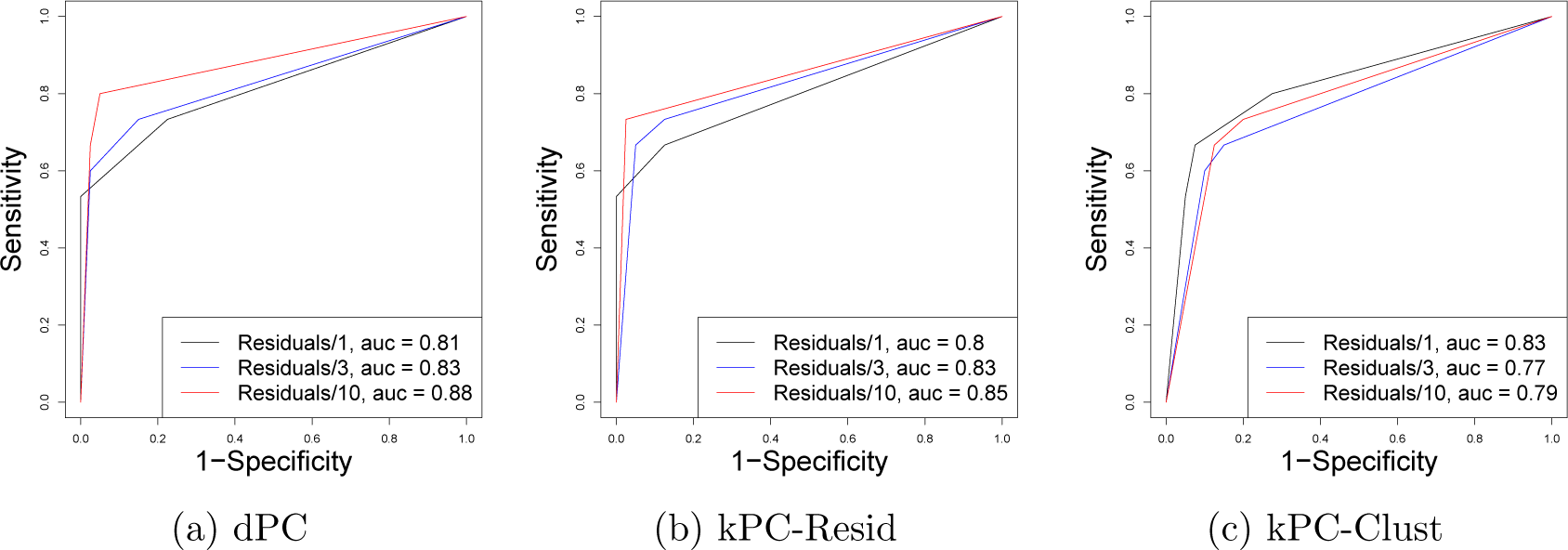
Signal to noise ratio effect on efficiency of the algorithms

Figure A3b in the supplementary material shows ROC curves for repeated simulations. The dPC, kPC-Resid, and kPC-Clust outperform the PC algorithm 99, 93, and 91 out of a 100 repetitions. To provide some insight into the variability of these results for different datasets, we show results for all simulated data from all 8 datasets in the supplementary material in Figure A5. Qualitatively the results are similar to that for dataset 7 presented here. In particular, the standard PC algorithm is almost always performing worst (except for dataset 2) and the dPC algorithm has a slight edge. Reducing noise helps improving results as shown in Figures A6, A7 and A8 for most of the dataset and algorithm combinations.

#### 3.2.3. Original data

Finally we tested all the algorithms on the original data from [20]. Again we only show the results on dataset 7 here, the rest of the results are provided in the supplementary material in Figure A9. We expect to find the skeleton of the graph in Figure 6a derived from Figure 1 as in the previous section. In Figure 6b we see the ROC curves for four versions of the PC algorithms. Results are very similar to the ones seen for simulated data: dPC and kPCs outperform PC and are quite similar among themselves, with dPC having a slight edge. We may conclude that independence criteria based PC versions are a significant improvement on the traditional PC algorithm on the real data as well as the simulated one.

**Figure 6.:**
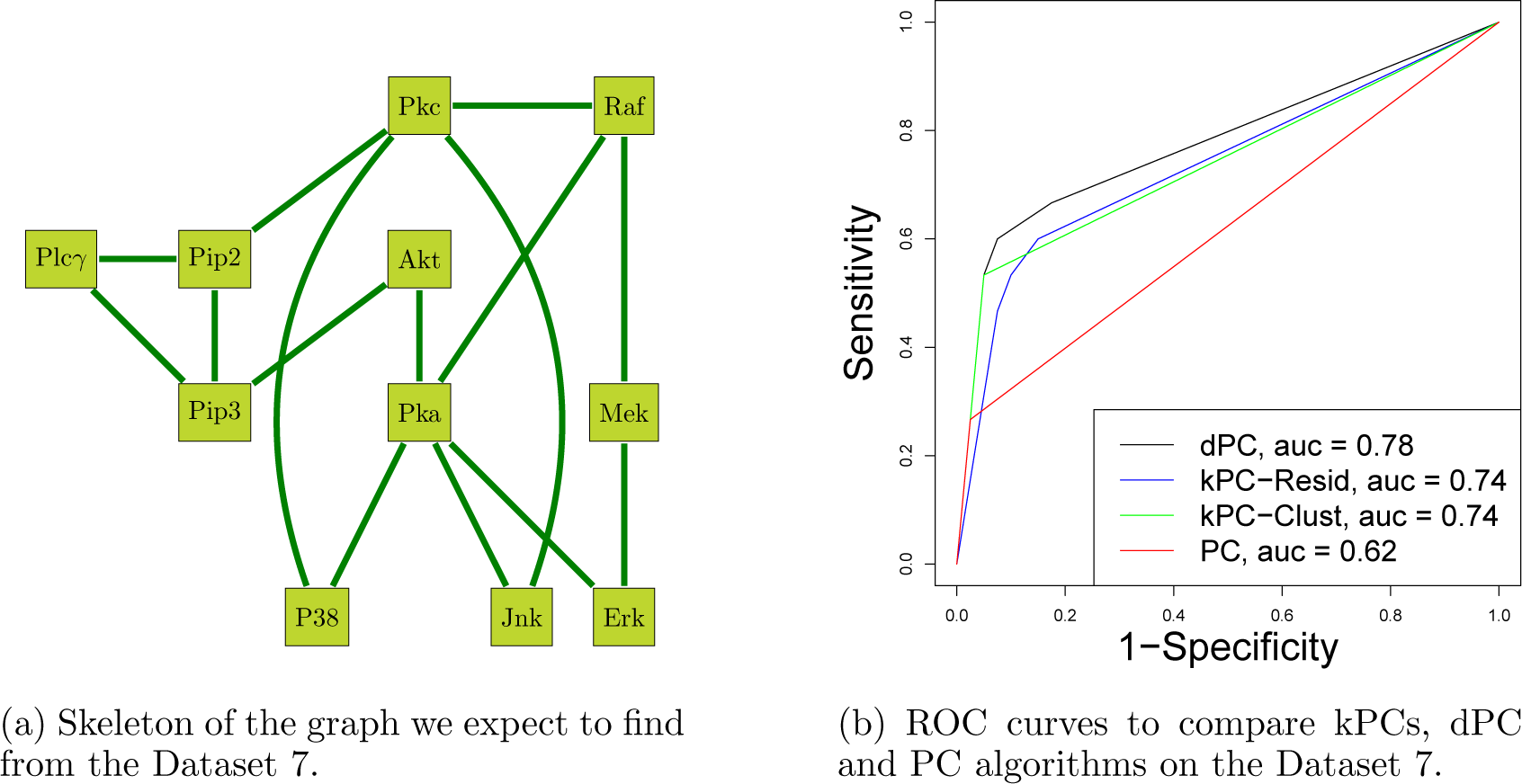
ROC curves for dataset 7.

### 3.3. Combining all datasets

The eight datasets of [20] have slightly different dependence structures due to variation in the external interventions. Combining information from all datasets should improve reconstruction of the underlying graph structure. To test this intuition we combine a consensus graphical structure from networks fitted to each dataset. Two types of consensus networks are obtained. The first takes edges that appear in *at least one* of the individual networks (labelled *union network*). The second calculates the typical average occurrence of edges over all edges in the union network and over all eight networks. Then only those edges of the union network are retained which occur (across all eight networks) more often than this typical average. We label this network *above-average network*. More sophisticated approaches are conceivable, however, here we only wanted to investigate whether there is potential improvement by combining networks at all, and the effect of the choice of an independence criterion on the consensus network. We compare the output of our algorithm to the skeleton illustrated in Figure 7a derived from Figure 1 and corresponding ROC curves are shown in Figure 7b and 7c.

**Figure 7.:**
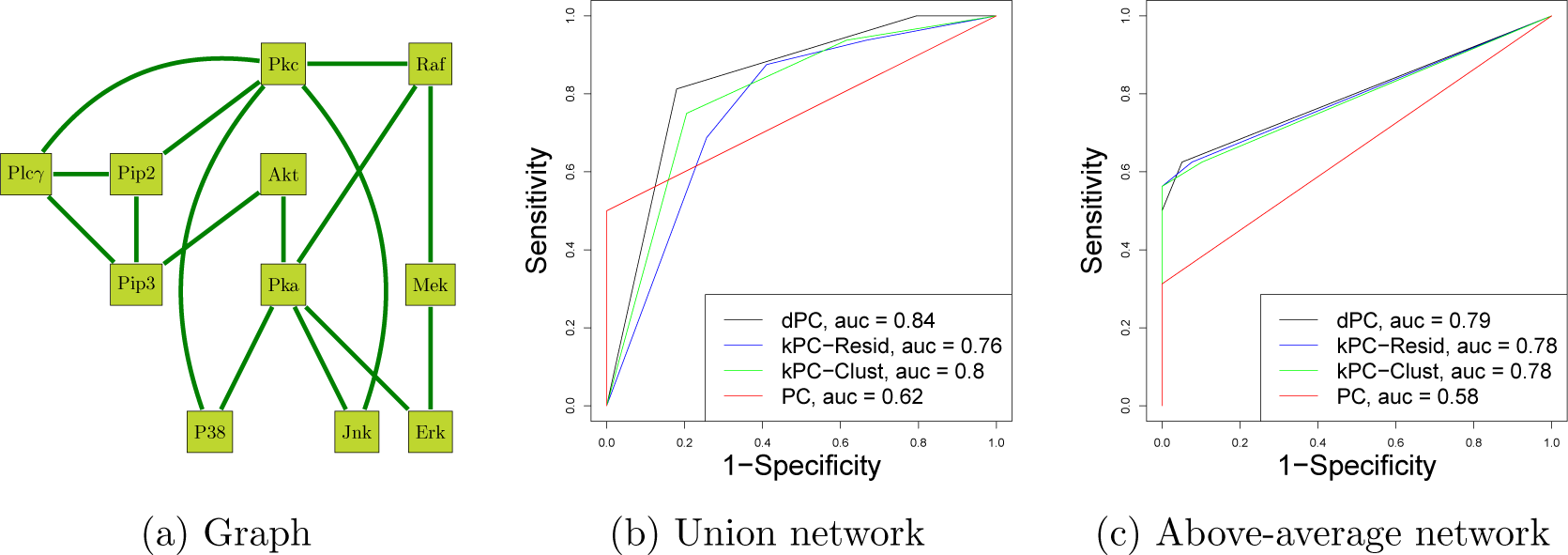
All 8 datasets combined.

Combining networks results in a slight improvement overall compared to Figure 6 with a trade-off between sensitivity and specificity shifted between the two types of combinations. The general independence criteria are again superior.

### 3.4. Discovering directions

So far we have looked at performance of algorithms when inferring the skeleton of a DAG with undirected edges only. The PC algorithm adds an edge orienting phase exploiting collider patterns and transitive closure requirements as formalised in the Meek rules [12] in the collider phase. Exploiting nonlinear relationships and non-Gaussian noise additional edges might be oriented. This is achieved by the PC algorithm extended by the generalised transitive phase which incorporates background knowledge emerging from testing directions exploiting nonlinearity and non-Gaussian noise.

With an imperfect independence oracle or data that do not strictly follow modelling assumptions, ambiguities can arise when orienting edges, possibly leading to cycles and doubly oriented edges. There are no general rules how to resolve such ambiguities. In this section we ignore doubly oriented edges as undirected for the purpose of assessing algorithms.

Results comparing algorithms by fractions of correct out of predicted orientations at various orientation phases are presented in Table 3. Free parameters were fixed as in the Table 2. The generalized transitive phase adds many more orientations to those found in the collider phase. For simulated data, this phase actually adds all the missing orientations in the correct direction.

**Table 3.:** The fraction of correct among predicted orientations at different stages of the algorithms.

|  | Experimental data |  |  |  | Simulated data |  |  |
| --- | --- | --- | --- | --- | --- | --- | --- |
|  | kPC Resid | kPC Clust | dPC |  | kPC Resid | kPC Clust | dPC |
| Collider/Transitive | 0 | 0 | 0 |  | 2/3 | 2/3 | 4/5 |
| Generalised transitive | 3/4 | 3/4 | 2/3 |  | 4/4 | 4/4 | 2/2 |
| Total | 3/4 | 3/4 | 2/3 |  | 6/7 | 6/7 | 6/7 |

We illustrate some of the results from Table 3. Figure 8 shows the output graphs of the orientation phases of algorithm kPC-Resid. There is one doubly oriented edge emerging in the collider phase between AKT *↔* PIP3. This phase also orients one edge in the wrong direction. Since there are no more colliders no further edge can be oriented at that stage. However, exploiting nonlinearity and non-Gaussian noise it is straightforward to orient the rest of the edges in the Generalised transitive phase.

**Figure 8.:**
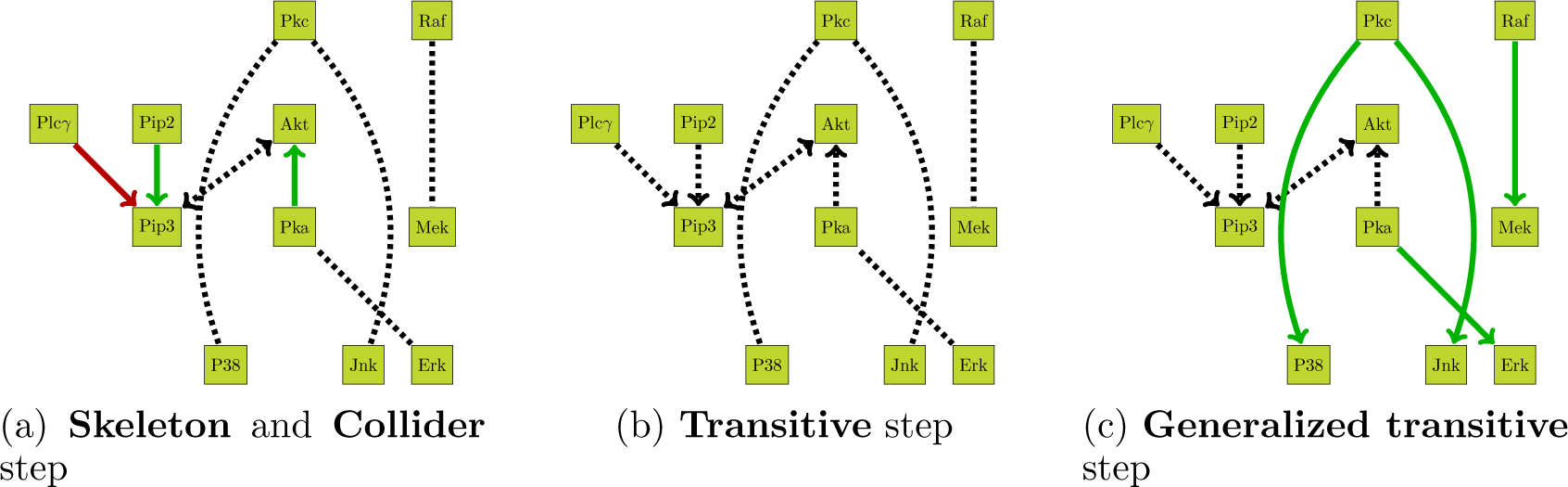
Output of the kPC-Resid algorithm on the data simulated from dataset 8. Color coding: dashed black undirected or doubly directed edges represent correctly identified undirected edges, green directed edges represent correct, while red directed edges represent incorrect orientations. Dashed black oriented edges are from the previous phase.

Similarly, Figure 9 shows the output of the kPC-Resid on the real dataset 8. Unfortunately the graph contains several triangles which make it impossible to orient any edge in the collider phase. The generalised transitive phase can differentiate between the oriented structures. Finally, for comparison with the kPC algorithm, Figure 10 shows the output of the dPC algorithm on simulated dataset 8. In contrast to the kPC-Resid algorithm there are enough colliders present to allow the algorithm to orient edges even in the collider phase. It orients four edges correctly, even though strictly speaking a collider at MEK is inconsistent with a collider at ERK. The rest of the edges gets oriented in the generalised transitive phase. Due to the two inconsistent colliders the overall result is slightly inferior to that of the kPC-Resid algorithm.

**Figure 9.:**
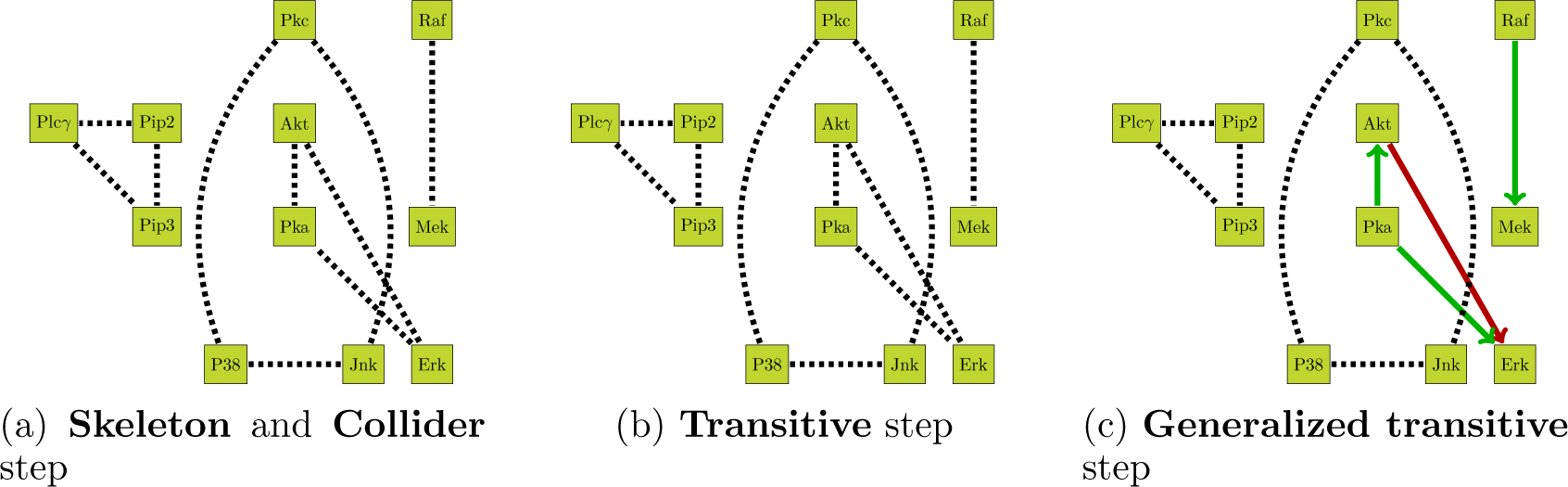
Output of the kPC-Resid algorithm on the dataset 8.

**Figure 10.:**
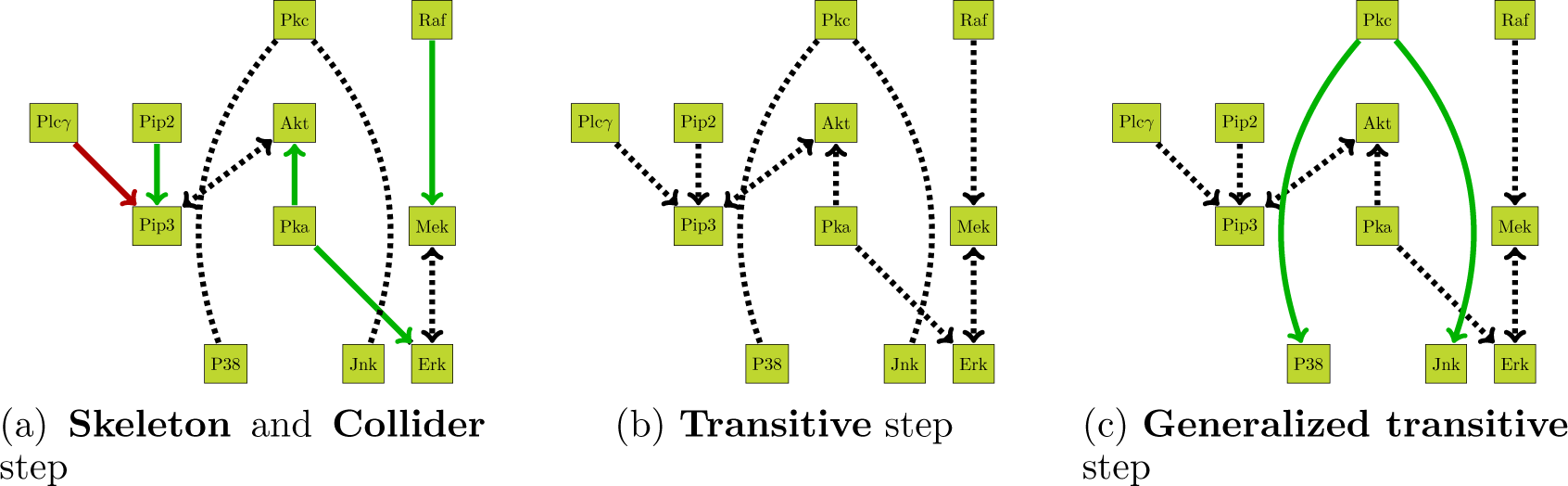
Output of the dPC algorithm on the data simulated from the dataset 8.

## 4. Discussion

The purpose of this study was to investigate how far probabilistic independence criteria for continuous data that go beyond linear relationships and Gaussian noise can improve the identification of edges and their orientation in a causal graph when applied to experimental data and data simulated in a realistic fashion from experimental data. We analysed two different criteria proposed in the literature, the Hilbert-Schmidt Independence Criterion or HSIC, and the Distance Covariance Criterion, or DCC in the context of the popular PC algorithm that relies on measures of probabilistic independence for network inference. The distance covariance is a natural extension of the Pearson correlation parameter [25]. To our knowledge this is the first implementation and application of the DCC to causal or network inference. All algorithms discussed in this study are available as package for the R statistical environment [18].

Overall, our findings confirm that the performance of general independence criteria is decisively better over that based on linear relationships with Gaussian noise on simulated as well as experimental data in terms of correct undirected edges as well as of correct directions. Secondly, we find only little difference between the performance of the HSIC and DCC in general, with the DCC showing slightly better performance for some datasets.

In order to assess the algorithms in a realistic scenario we applied them to a wellknown experimental dataset for which the network is approximately known based on biological knowledge. Of course, this knowledge of the network might be inaccurate and we therefore propose a generic way to simulate data based on experimental data and an approximate or putative network structure that keeps much of the noise characteristics of the original data but reproduces those and only those conditional dependencies required by the network. As we demonstrate in our analysis these simulated datasets form an excellent compromise between retaining much of the nonlinear and non-Gaussian characteristics of the original data, but for an exactly known network. As we see in our study one difficulty remains: if some arcs of the assumed network are not supported by the data, for example, if there is little dependency in the data in the first place between two variables which we wish to connect in the network, our method is unable to create such dependency artificially. Nevertheless, as long as the assumed network reflects most of the dependencies in the experimental data, the simulated data are useful for comparative studies between different algorithms as shown in section 3.2.2.

The PC algorithm requires a test for conditional independence. Independence criteria might, however, only be available in an unconditional form. We propose a simple procedure, based on fitting nonlinear regressions, to adapt such criteria to the conditional independence case. Since there is a conditional version of the HSIC available we had an opportunity to compare the conditional HSIC (in algorithm kPC-Clust) with our adaptation of the unconditional HSIC (in algorithm kPC-Resid). As can be seen throughout the study, the adapted kPC-Resid version, particularly on experimental and realistically simulated data, is performing comparably to the conditional version with the conditional HSIC having a slight advantage. It is worth noting that kPC-Clust involves calculating the empirical estimate of the conditional HSIC (3) which is computationally significantly more expensive than the unconditional HSIC (2), therefore in practice kPC-Clust can be up to 5 *−* 10 times slower than kPC-Resid or dPC.

The empirical estimation of HSIC depends on parameters such as the kernel width *λ* and the regularisation parameter *E*. Here we used simulations to find sensible ranges for these parameters. Another advantage of the DCC is that it is less affected by parameter choices, essentially only the power parameter *ξ* which can be safely set to a value between 0.1 and 1 without affecting the results too much.

The PC algorithm is very restrictive in its assumptions on the dependency structure. For example, cycles or unobserved variables are excluded. It would be interesting to see whether inference techniques allowing such more complex assumptions benefit from general independence criteria in the same way the PC algorithm does.

The PC algorithm is firmly based in a frequentist statistical framework. Bayesian inference is often strongly dependent on specific noise models through the likelihood function. It needs to be explored how to incorporate independence criteria in a Bayesian framework, possibly through a form of loss likelihood [1].

## Appendix A. Supplementary Material

**Table A1.:**
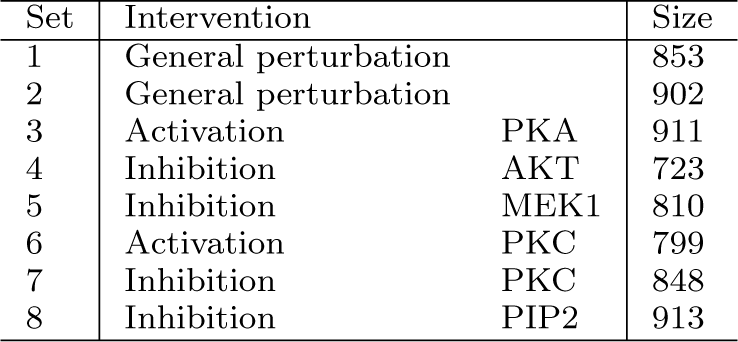
Table of datasets

**Figure A1.:**
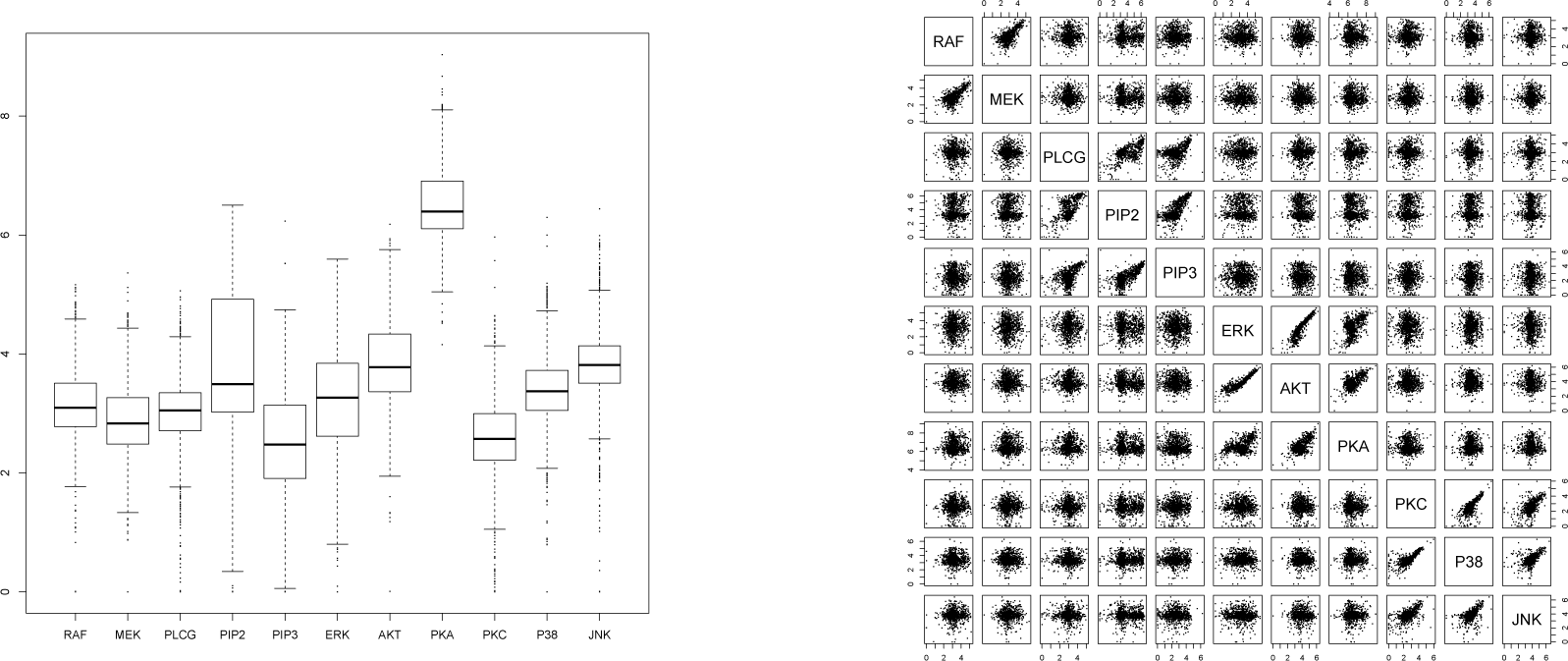
Boxplot and pair plot of dataset 8 after log transformation.

**Figure A2.:**
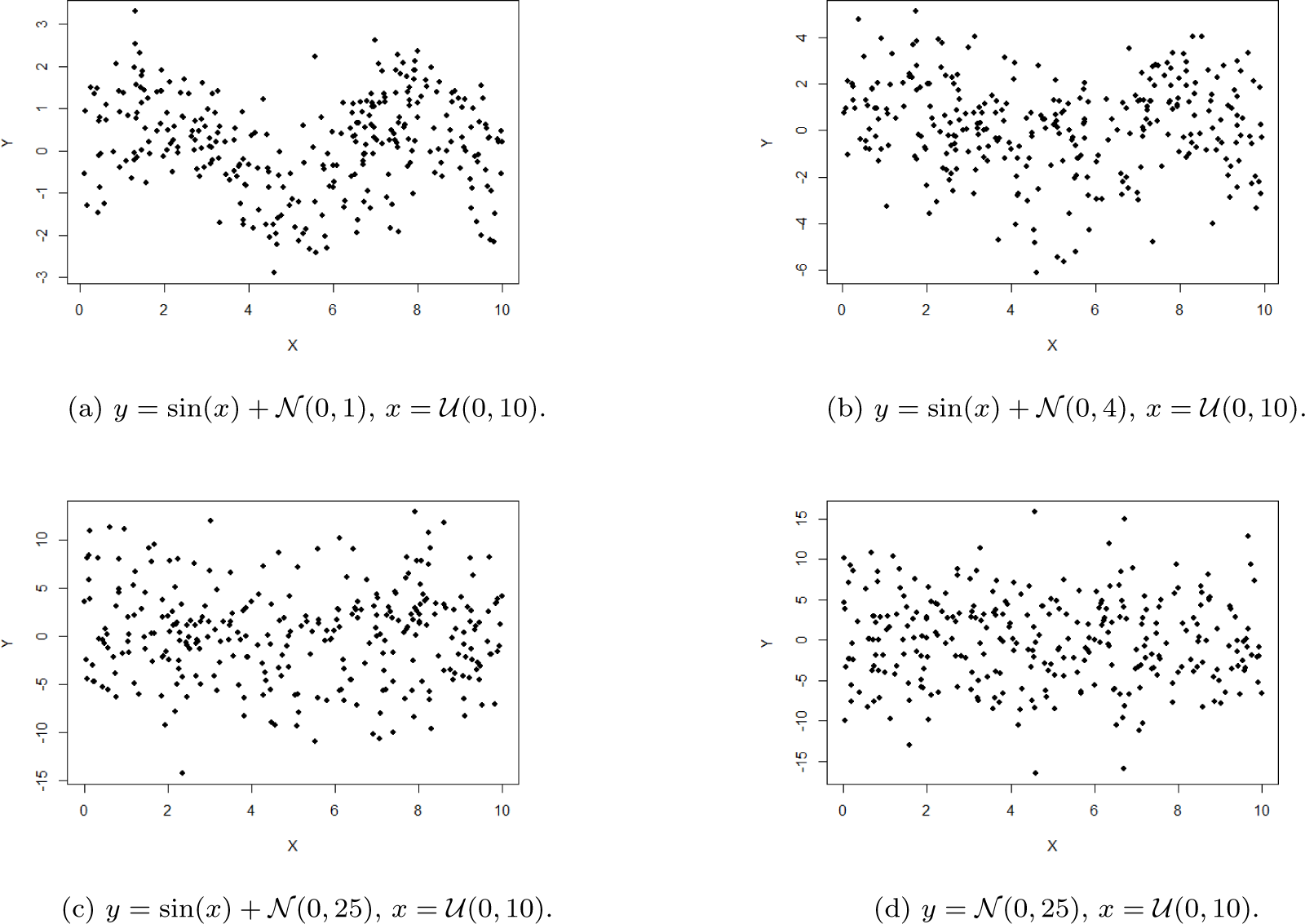
Data simulated with nonlinear dependencies

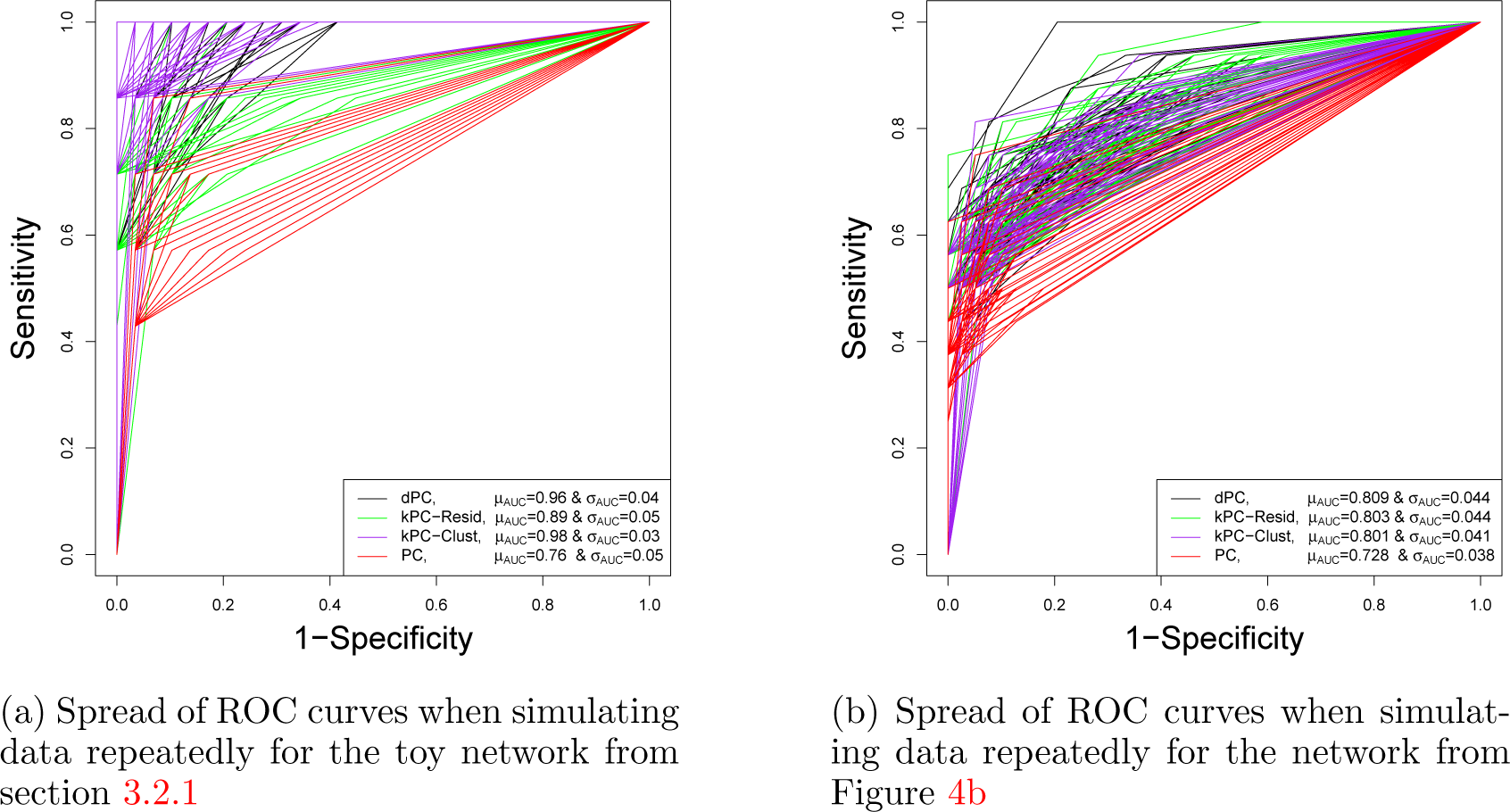

**Figure A4.:**
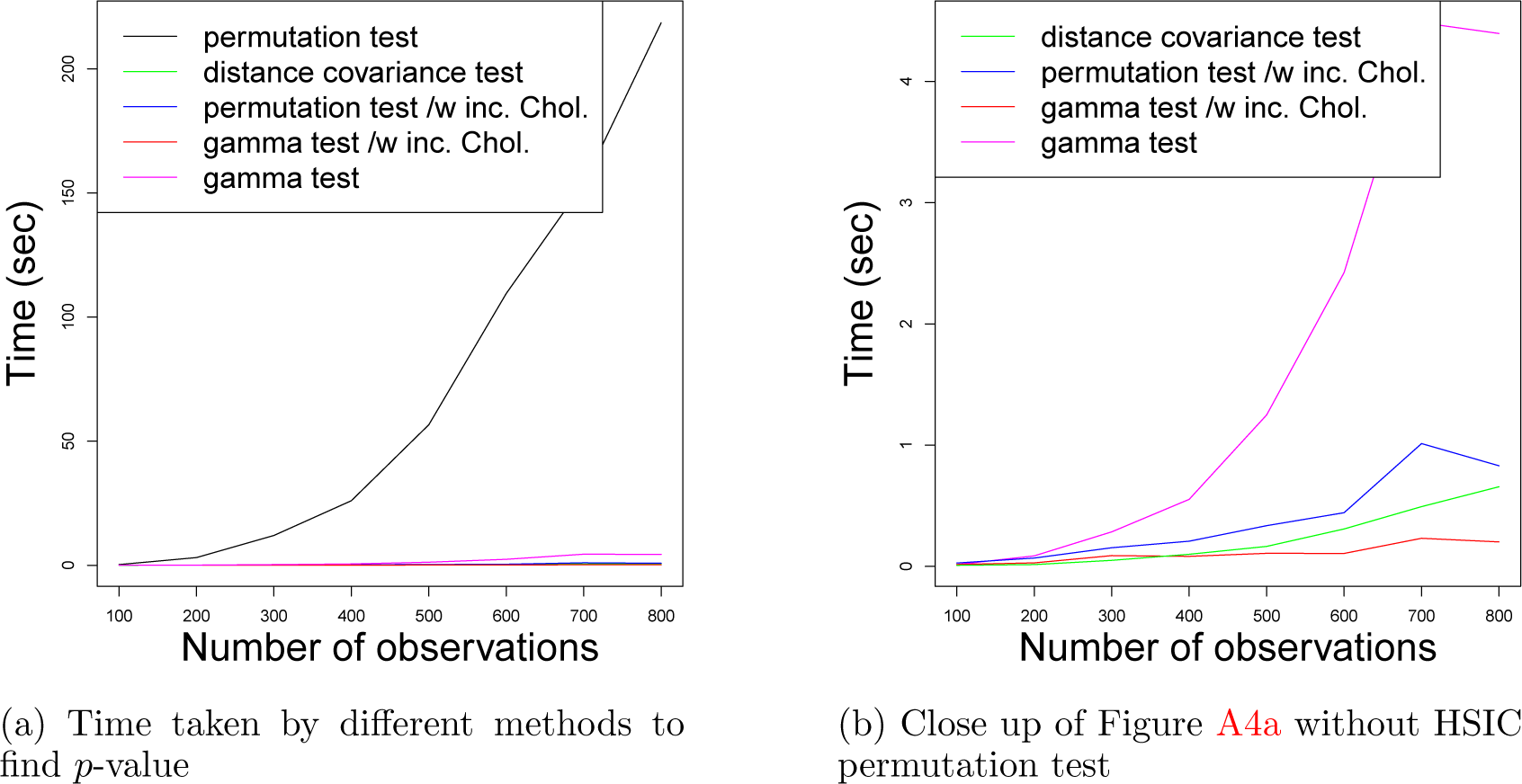
Time efficiency of independence tests.

**Figure A5.:**
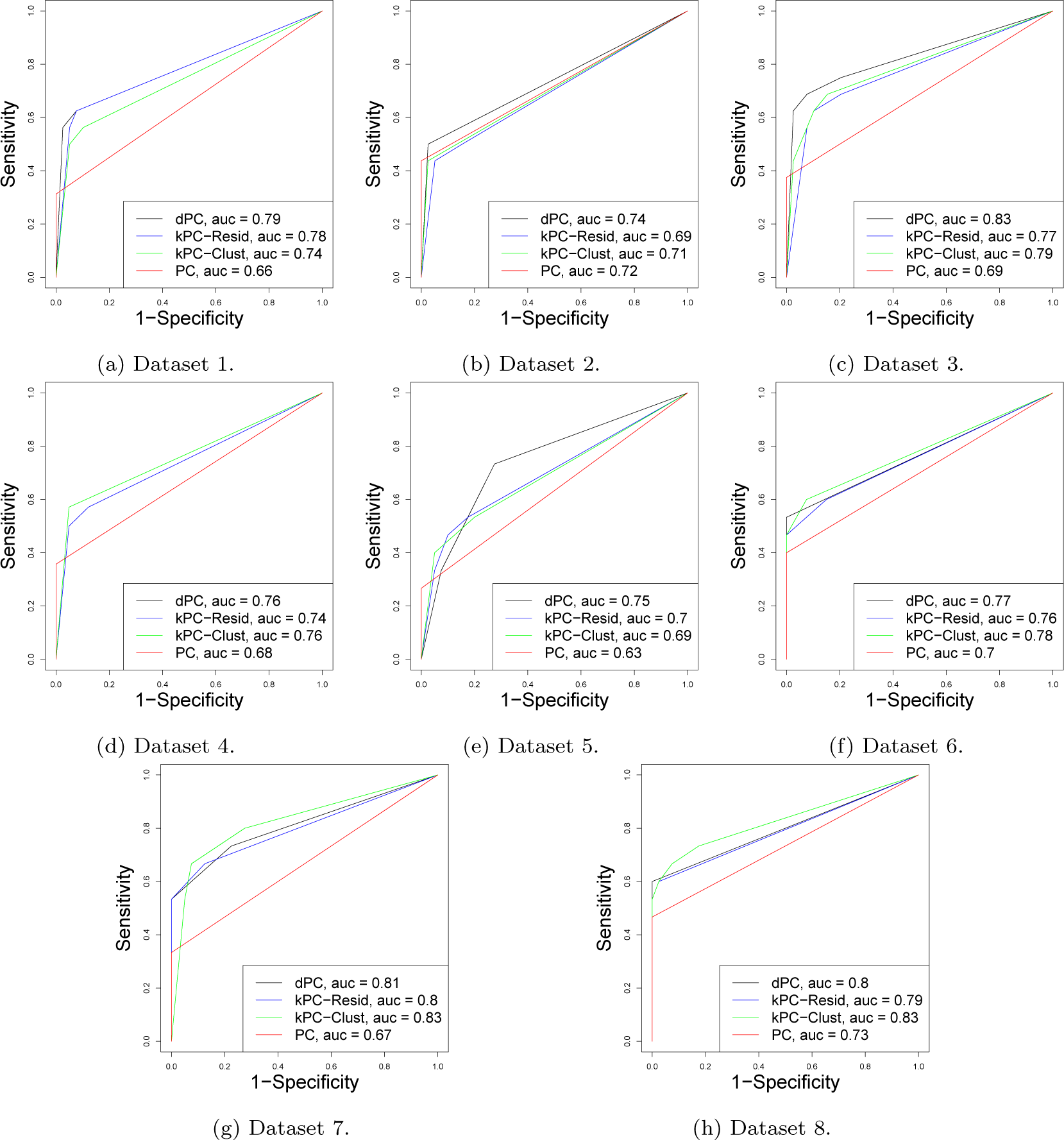
ROC curves to compare kPCs, dPC and PC algorithms on data simulated from the indicated datasets.

**Figure A6.:**
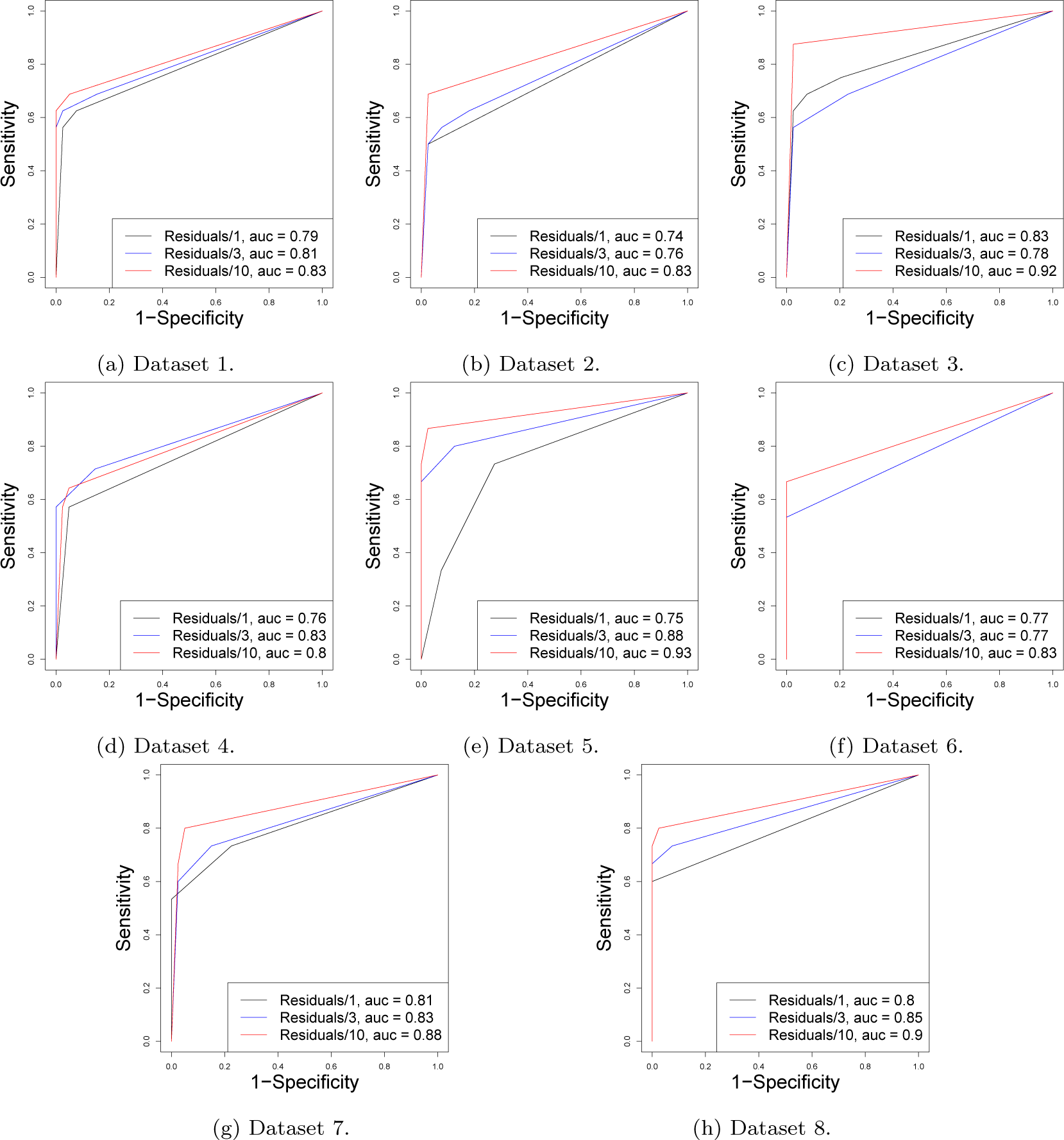
dPC effectiveness on data with varying noise levels, all eight datasets.

**Figure A7.:**
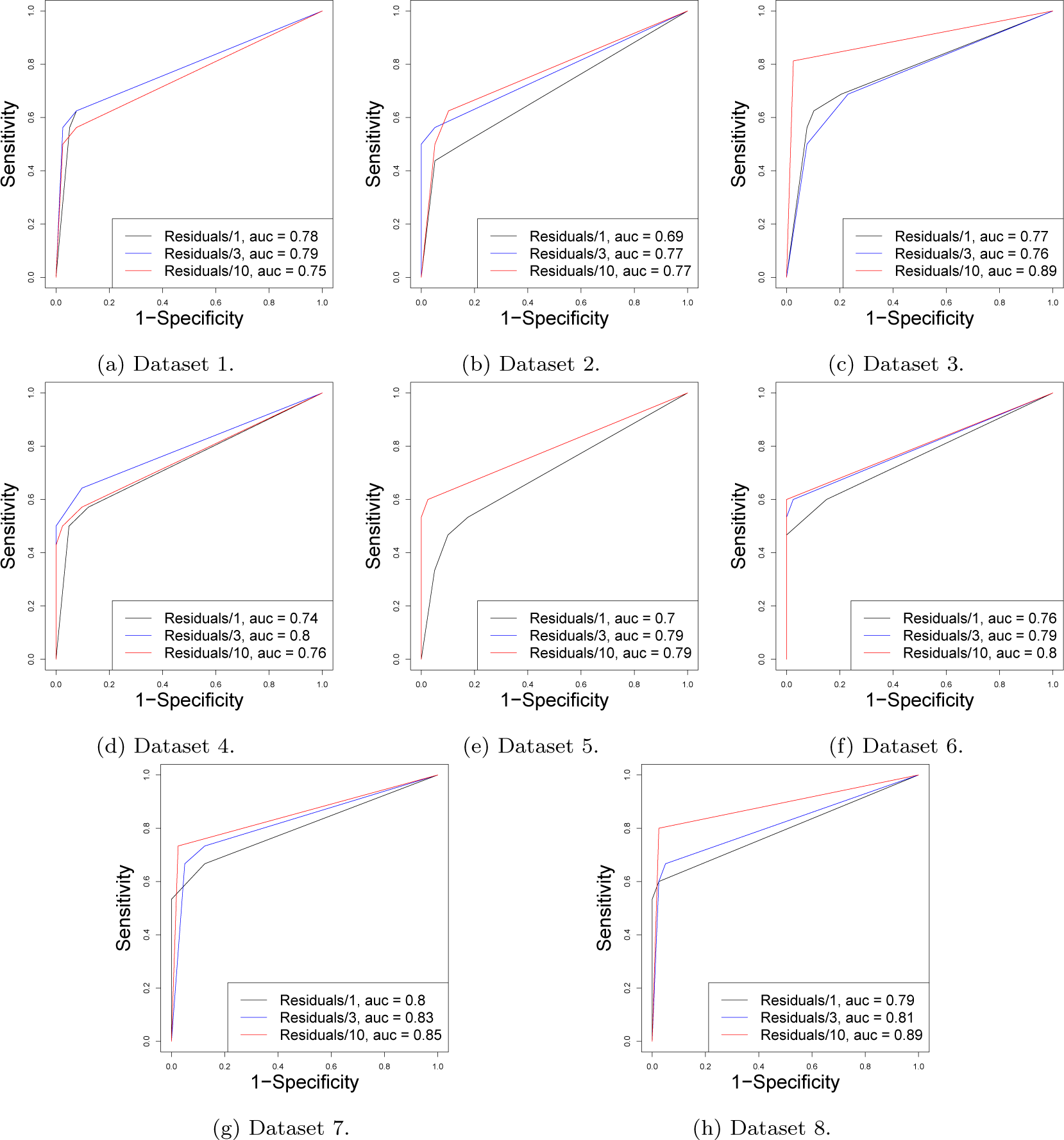
kPC-Resid effectiveness on data with varying noise levels, all eight datasets.

**Figure A8.:**
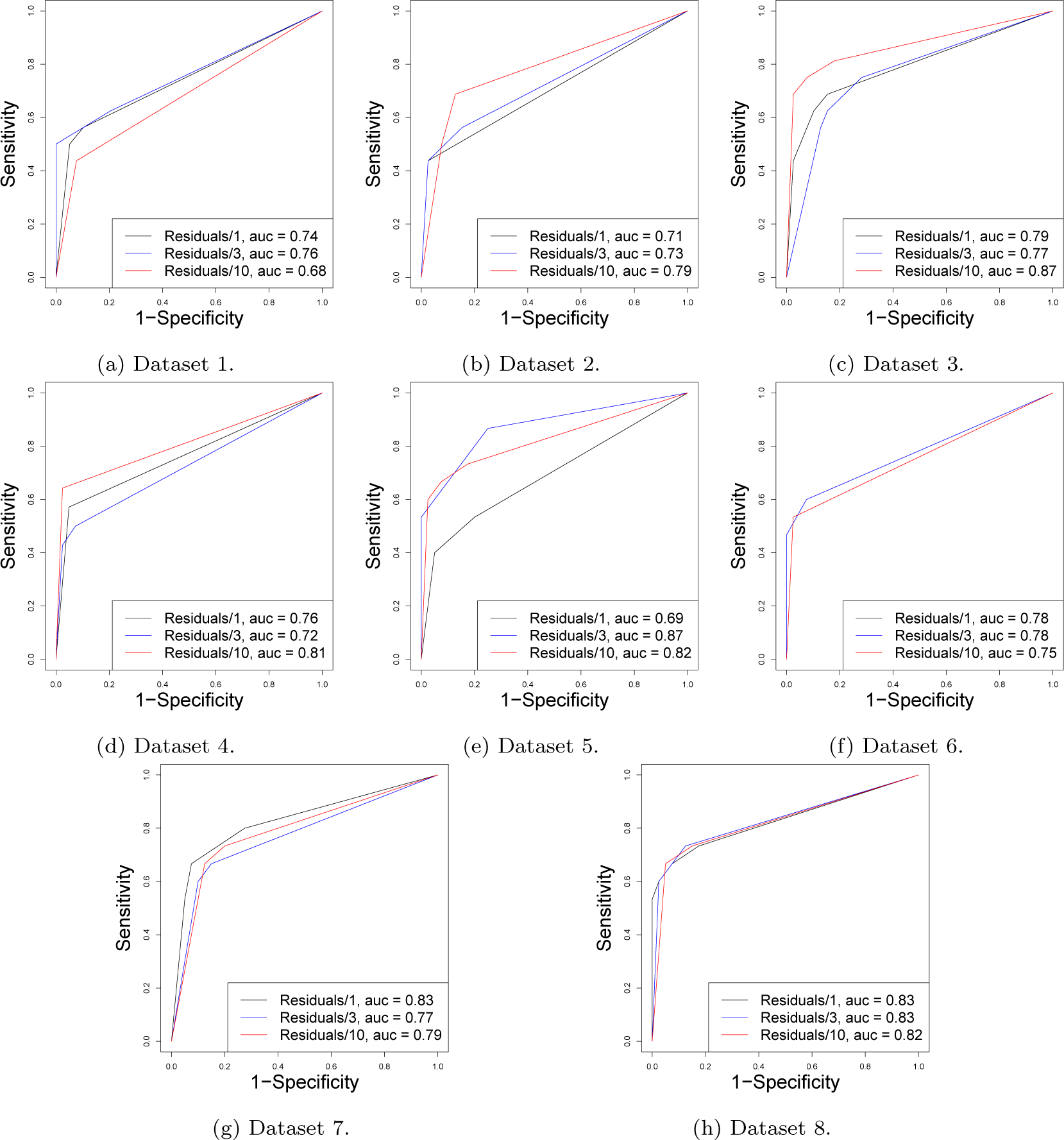
kPC-Clust effectiveness on data with varying noise levels, all eight datasets.

**Figure A9.:**
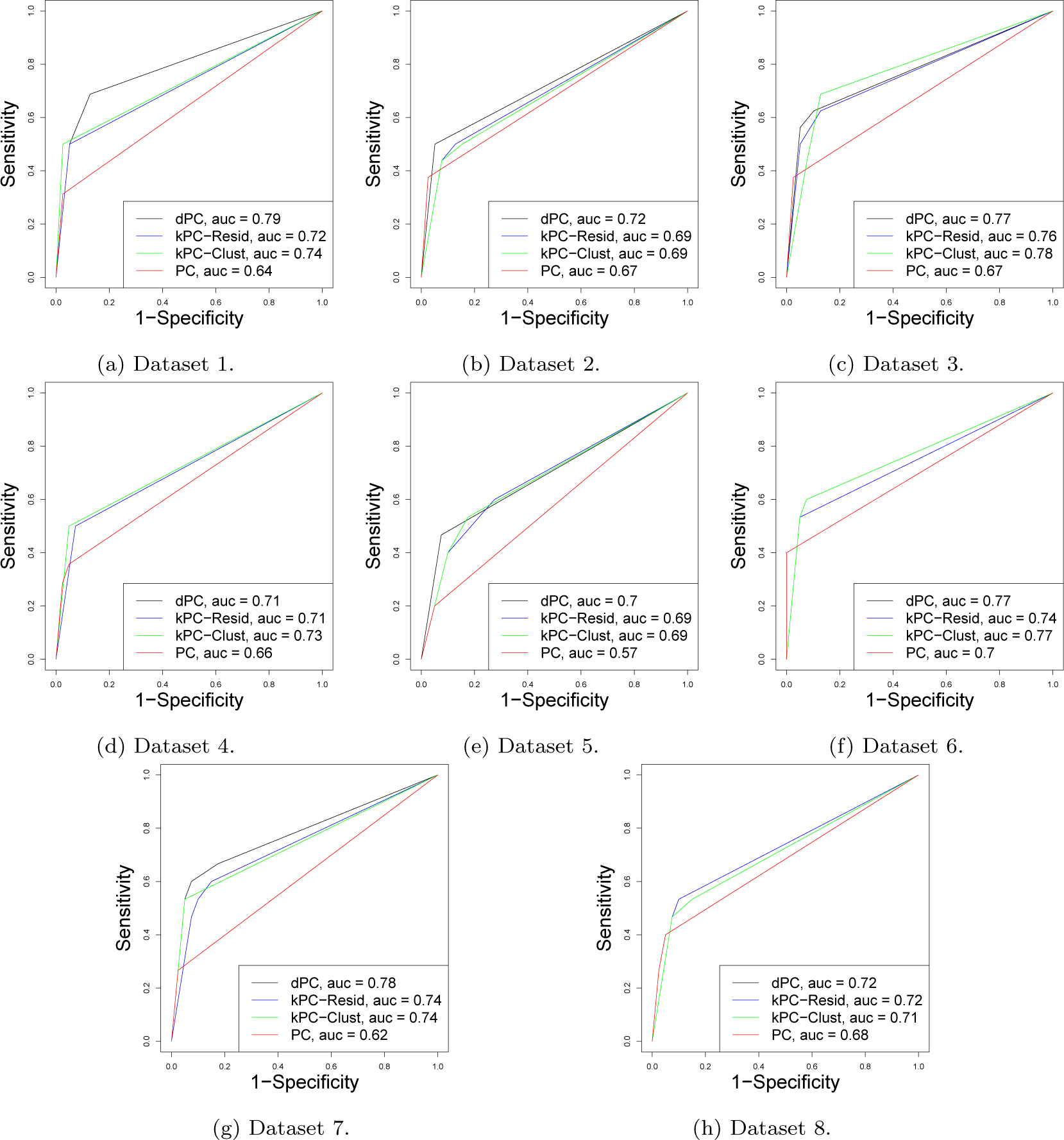
ROC curves to compare kPCs, dPC and PC algorithms on the real data.

## References

[1] P. Bissiri, C. Holmes, and S.G. Walker, A general framework for updating belief distributions, Journal of the Royal Statistical Society: Series B (Statistical Methodology) (2016).

[2] A.J. Butte and I.S. Kohane, Mutual information relevance networks: functional genomic clustering using pairwise entropy measurements, in Pac Symp Biocomput, Vol. 5. 2000, pp. 418–429.

[3] J.J. Faith, B. Hayete, J.T. Thaden, I. Mogno, J. Wierzbowski, G. Cottarel, S. Kasif, J.J. Collins, and T.S. Gardner, Large-scale mapping and validation of escherichia coli transcriptional regulation from a compendium of expression profiles, PLoS biol 5 (2007), p. e8.

[4] A. Gretton, O. Bousquet, A. Smola, and B. Schölkopf, Measuring statistical dependence with Hilbert-Schmidt norms, in Algorithmic learning theory. Springer, 2005, pp. 63–77.

[5] A. Gretton, K. Fukumizu, C.H. Teo, L. Song, B. Schölkopf, and A.J. Smola, A kernel statistical test of independence, NIPS20 (2008).

[6] J.A. Hartigan and M.A. Wong, A k-means clustering algorithm., Applied Statistics 28 (1979), pp. 100–108.

[7] T. Hastie and R. Tibshirani, Generalized additive models, Statistical science (1986), pp. 297–310.

[8] P.O. Hoyer, D. Janzing, J.M. Mooij, J. Peters, and B. Schölkopf, Nonlinear causal discovery with additive noise models, in Advances in neural information processing systems. 2009, pp. 689–696.

[9] A. Irrthum, L. Wehenkel, P. Geurts, et al., Inferring regulatory networks from expression data using tree-based methods, PloS one 5 (2010), p. e12776.

[10] C. Lippert, O. Stegle, Z. Ghahramani, and K. Borgwardt, A kernel method for unsupervised structured network inference, JMLR Workshop and Conference Proceedings Volume 5:368–375 (2009).

[11] D. Marbach, J.C.C. and Robert Kffner, N.M. Vega, R.J. Prill, D.M. Camacho, K.R.A. amd The DREAM5 Consortium, M. Kellis, J.J. Collins, and G. Stolovitzky, Wisdom of crowds for robust gene network inference, Nature Methods 9, 796–804 (2012) (2012).

[12] C. Meek, Causal inference and causal explanation with background knowledge, in Proceedings of the Eleventh conference on Uncertainty in artificial intelligence. Morgan Kaufmann Publishers Inc., 1995, pp. 403–410.

[13] P.E. Meyer, K. Kontos, F. Lafitte, and G. Bontempi, Information-theoretic inference of large transcriptional regulatory networks, EURASIP journal on bioinformatics and systems biology 2007 (2007), pp. 1–9.

[14] T. Obayashi and K. Kinoshita, Rank of correlation coefficient as a comparable measure for biological significance of gene coexpression, DNA research 16 (2009), pp. 249–260.

[15] R. Opgen-Rhein and K. Strimmer, From correlation to causation networks: a simple approximate learning algorithm and its application to high-dimensional plant gene expression data, BMC systems biology 1 (2007), p. 37.

[16] J. Pearl, Causality, Cambridge university press, 2009.

[17] R Core Team, R: A Language and Environment for Statistical Computing, R Foundation for Statistical Computing, Vienna, Austria. Available at http://www.R-project.org.

[18] R Core Team, cR: A Language and Environment for Statistical Computing, R Foundation for Statistical Computing, Vienna, Austria (2014). Available at http://www.R-project.org/.

[19] M.L. Rizzo and G.J. Szekely, E-statistics (2014). Available at http://cran.r-project.org/web/packages/energy/energy.pdf.

[20] K. Sachs, O. Perez, D. Pe’er, D.A. Lauffenburger, and G.P. Nolan, Causal protein-signaling networks derived from multiparameter single-cell data, Science 308 (2005), pp. 523–529.

[21] D. Sejdinovic, B. Sriperumbudur, A. Gretton, K. Fukumizu, et al., Equivalence of distance-based and rkhs-based statistics in hypothesis testing, The Annals of Statistics 41 (2013), pp. 2263–2291.

[22] J. Shawe-Taylor and N. Cristianini, Kernel methods for pattern analysis, Cambridge university press, 2004.

[23] P. Spirtes, C.N. Glymour, and R. Scheines, Causation, prediction, and search, Vol. 81, MIT press, 2000.

[24] G.J. Székely, M.L. Rizzo, N.K. Bakirov, et al., Measuring and testing dependence by correlation of distances, The Annals of Statistics 35 (2007), pp. 2769–2794.

[25] G.J. Székely, M.L. Rizzo, et al., Brownian distance covariance, The annals of applied statistics 3 (2009), pp. 1236–1265.

[26] R.E. Tillman, A. Gretton, and P. Spirtes, Nonlinear directed acyclic structure learning with weakly additive noise model, NIPS 22, Vancouver (2009).

[27] S. Wood, Stable and efficient multiple smoothing parameter estimation for generalized additive models., Journal of the American Statistical Association. 99 (2004), pp. 673–686.

[28] S. Wood, Fast stable restricted maximum likelihood and marginal likelihood estimation of semiparametric generalized linear models., Journal of the Royal Statistical Society. 73 (2011), pp. 3–36.

